# AIM (Angular Indication Measurement)- Visual Acuity: An adaptive, self-administered, and generalizable vision assessment method used to measure visual acuity

**DOI:** 10.1101/2023.02.25.529586

**Authors:** Jan Skerswetat, Jingyi He, Jay Bijesh Shah, Nicolas Aycardi, Michelle Freeman, Peter John Bex

## Abstract

This proof-of-concept study introduces Angular Indication Measurement and applies it to VA (AIM-VA). First, we compared the ability of AIM-VA and ETDRS to detect defocus and astigmatic blur in 22 normally-sighted adults. Spherical and cylindrical lenses (±0.00D, +0.25D, +0.50D, +0.75D, +1.00D, +2.00D and +0.50D, +1.00D, +2.00D each at 0°, 90°, 135°, respectively) in the dominant eye induced blur. Second, we compared repeatability over two tests of AIM-VA and ETDRS. A 2-way-ANOVA showed a main effect for defocus-blur and test with no interaction. A 3-way-ANOVA for the astigmatism experiment revealed main effects for test type, blur, and direction and with no interactions. Planned multiple comparisons showed AIM had greater astigmatic-induced VA loss than ETDRS. Bland-Altman plots showed small bias and no systematic learning effect for either test type and improved repeatability with >2 adaptive steps for AIM-VA. AIM-VA’s ability to detect defocus was comparable with that of an ETDRS letter chart and showed greater sensitivity to astigmatic blur, and AIM-VA’s repeatability is comparable with ETDRS when using 2 or more adaptive steps. AIM’s self-administered orientation judgment approach is generalizable to interrogate other visual functions, e.g., contrast, color, motion, stereo-vision.

## 1. Introduction

Visual function assessment is an essential task for eye care clinicians. Visual acuity (VA) is arguably the key endpoint in current clinical practice to assess central vision as it is sensitive to changes of the refraction of the eye (1), but also changes of central retinal (2) and ocular media structures (3), changes along the visual pathway (4), and atypical visual-cortical structures (5). Since the first known reports for VA tests by Tobias Mayer 1757 (6) and Heinrich Küchler in 1843 (7), various techniques and tasks have evolved to measure VA; however, all of those tests examine the observer’s ability to discern different structural details from each other by subjective report, for recent review see (6).

One of the major challenges for eyecare professionals worldwide is the accurate correction of refractive errors, where estimates suggest that approximately 2.6 billion people are affected by myopia and 1.8 billion people by presbyopia (8). Defocus blur, i.e., isotopically distributed blur, may reduce VA in cases of myopia and higher hypermetropia and can be corrected with spherical lenses. Astigmatism, i.e. anisotopic blur with two focal lines of different focal lengths, causes uneven defocus and may generate systematic distortions of an image on the retina.

In practice, VA assessment typically requires a participant to identify letter or number optotypes in 4 alternative forced choice (AFC) letter charts such as HOTV (9), 10AFC letter charts such as Early Treatment of Diabetic Retinopathy Study (ETDRS) (10, 11) or 9AFC letter charts such as Snellen (12). Observers who are non-literate or unfamiliar with the Roman alphabet may be asked to identify the orientation of the gap of tumbling E (13) or Landolt C (14) optotypes in 4 or 8 AFC tasks; to match child-friendly optotypes such as Lea symbols (15) or Auckland Optotypes (16) in 10AFC tasks; or to report the offset of misaligned line stimuli in 2AFC tasks (17). For participants such as babies who are unable to generate a verbal or manual report, preferential looking behavior can be converted to a 2AFC task (18).

Typically, optotypes are presented in printed or computer-generated charts in which the layout, size and spacing of optotypes is organized with monotonically increasing difficulty. The number and spacing of letters can vary (12) or may be logarithmic (19) following the Weber-Fechner law (20). Various rules have been developed to score the correct and incorrect responses of the participant, including letter scoring, in which VA is related to the total number of letters correctly identified; line scoring, in which VA is defined by the optotype size where a specified proportion of letters are correctly reported; and stopping criteria, all of which affect the estimate of VA and repeatability (21–23).

The transition to computer-administered tests avoids memorization artifacts of printed charts and allows the implementation of adaptive psychophysical testing principles (24, 25) and automated scoring eliminating inter-examiner differences (24, 26). More recently, numerous web-browser- and app-based VA tests have entered the field to monitor VA remotely (25, 27, 28).

Current tests do not capture an individual’s performance change with changing visibility levels for targets of differing size but rather only report the threshold, or acuity at a single point estimated from the psychometric function. The range between reliably detecting an optotype or its orientation and plateauing in performance is currently not captured, however parameters such as the slope of the psychometric function may be informative as well as they can be affected by blur (29). Also, an individual’s baseline error rate (i.e., noise or uncertainty for supra-threshold letters that are correctly identified) is not captured using chart-based VA tests and might be systematically disrupted due to factors such as visual distortions.

To address the shortcomings of current visual acuity tests described above (namely optotype structure, familiarity and # alternatives; behavioral task; and data scoring and fitting), we introduce the Angular Indication Measurement (AIM) method, which overcomes these issues and apply it to measure VA. AIM is computer-based, generalizable, self-administered, and response-adaptive. AIM can be applied to measure multiple visual function endpoints by adapting different stimulus and response formats to an orientation judgment. In each case, stimuli are presented at a range of personalized stimulus levels and random orientations (acuity, color, contrast, stereoscopic disparity etc.) or directions (motion). The observer’s task is to report the perceived orientation or direction of the stimulus. The difference between the actual and reported stimulus orientation/direction provides a continuous error score as a function of stimulus intensity (30). For VA, observers report the orientation of randomly-oriented C-like stimuli of varying sizes, which requires no literacy or memory for optotypes by the participant and automated data analysis for the examiner.

The current study has three objectives. First, we introduce AIM for Visual Acuity assessment (AIM-VA) and describe its features and outcome measures. Second, we report AIM-VA’s ability to detect defocus and astigmatic blur induced with ophthalmic lenses in normally-sighted adults and compare it with results of a conventional ETDRS letter chart (Experiment 1). Third, we investigate the repeatability of AIM-VA and compare it with ETDRS results (Experiment 2).

## 2. Methods

Participants were recruited from the translational vision laboratory and from the undergraduate population at Northeastern University, Boston. Undergraduate students received course credit towards the completion of their Introductory Psychology course in exchange for their participation. Written and verbal information about the project were provided in advance to the participants and they gave informed consent via online form before taking part. The Institutional Review Board of Northeastern University approved the study and was in line with the ethical principles of the Helsinki declaration of 1975.

### 2.1 Participants

#### Pilot experiment

Eleven adults from Northeastern University’s student cohort without a known (self-reported) ocular opacity, accommodative or extreme refractive disorder/disease took part.

#### Experiment 1

Twenty-three adults from Northeastern University’s student cohort and members of the Translational Vision Science lab (including JS, MF, JBS, JH) took part in the study. One participant was excluded due to erroneously duplicated data. Of the remaining twenty-two participants (age range 18-35 years), eight wore no optical correction, fourteen wore correction. An objective refraction was performed using autorefractometer (AR(K)7610 Auto Refracto-Keratometer) and prescription of glasses if worn, were measured using an auto-focimeter (Xinyuan, Auto Lensmeter). Although no data on subjective, optometrist-performed refraction was available, we assume that participants who wore corrections were best-corrected.

#### Experiment 2

Participants from experiment 1 participated also in the second experiment.

#### Control experiment

An additional control experiment was performed to determine the effect of the number of adaptive steps on repeatability, twelve participants were recruited, including a subset of seven participants from Experiment 2 and five additional participants (three with and two without correction).

### 2.2 Equipment

Stimuli were presented on a HP (USA) All-in-One computer and monitor system (Model: 32-a1027c Rfrbd PC) with a framerate of 60 Hz with a pixel resolution of [3840 / 2160], that subtended 10° at a 4m viewing distance with a resolution of 384 pixels per degree. Stimuli were generated using Matlab (MathWorks, R2021a) software in combination with *The Psychophysics Toolbox* (31, 32). An HP Bluetooth mouse was used as indication device. The monitor was gamma-corrected using datacolor Spyder (Switzerland) photometer. We used a retro-luminant ETRDS chart (Lighthouse, USA, second edition) and measured the background luminance with a LS-110 (Konica Minolta Sensing Americas) photometer.

### 2.3 AIM principal

AIM is a generalizable psychophysical paradigm in which one or more stimuli is presented at a random orientation, and the observer’s task is to indicate on a surrounding response ring the perceived orientation of the target, using a mouse or touch response. The difference between the true and reported orientation provides a continuous error score for each stimulus test level/intensity. Across different tests, stimulus intensity can refer to contrasts and spatial frequencies (33), cone-isolated contrasts and color differences (34), stereoscopic disparities (30) etc. In the current study, we varied optotype size to estimate VA. AIM combines a number of features that address current problems of other tests and also provides additional information about an observer’s performance, namely a) generalizable orientation judgment method, b) inclusive method as AIM is easy to perform due to the simple task nor does it require knowledge of letters or numbers, c) self-administered, thus reducing effort on part of the investigator, d) response-adaptive, thus narrowing in on individual threshold, e) continuous data, enabling personalized modeling of visual performance, f) active engagement of the user due to the search aspect of the task, g) instant analysis due to computerized approach, h) digital, thus deployable on a wide range of computerized screens, e.g. phones, tablets, laptops, computer monitors.

We used C optotypes as they have the advantage that the physical and spectral properties of optotypes do not vary within a chart except by the gap orientation and do not vary in meaning, unlike object optotypes such as letters, hence the acuity determined with such stimuli is a more unbiased estimate of an observer’s VA. Following International Council of Ophthalmology (30) recommendations, Cs were defined as interrupted circles whose stroke width and gap width are one-fifth of its outer diameter. The only difference of our C stimulus and a Landolt C is that the inner diameter had a smaller width than the outer part as an arc was used to construct the gap (Figure 1B).

**Figure 1:**
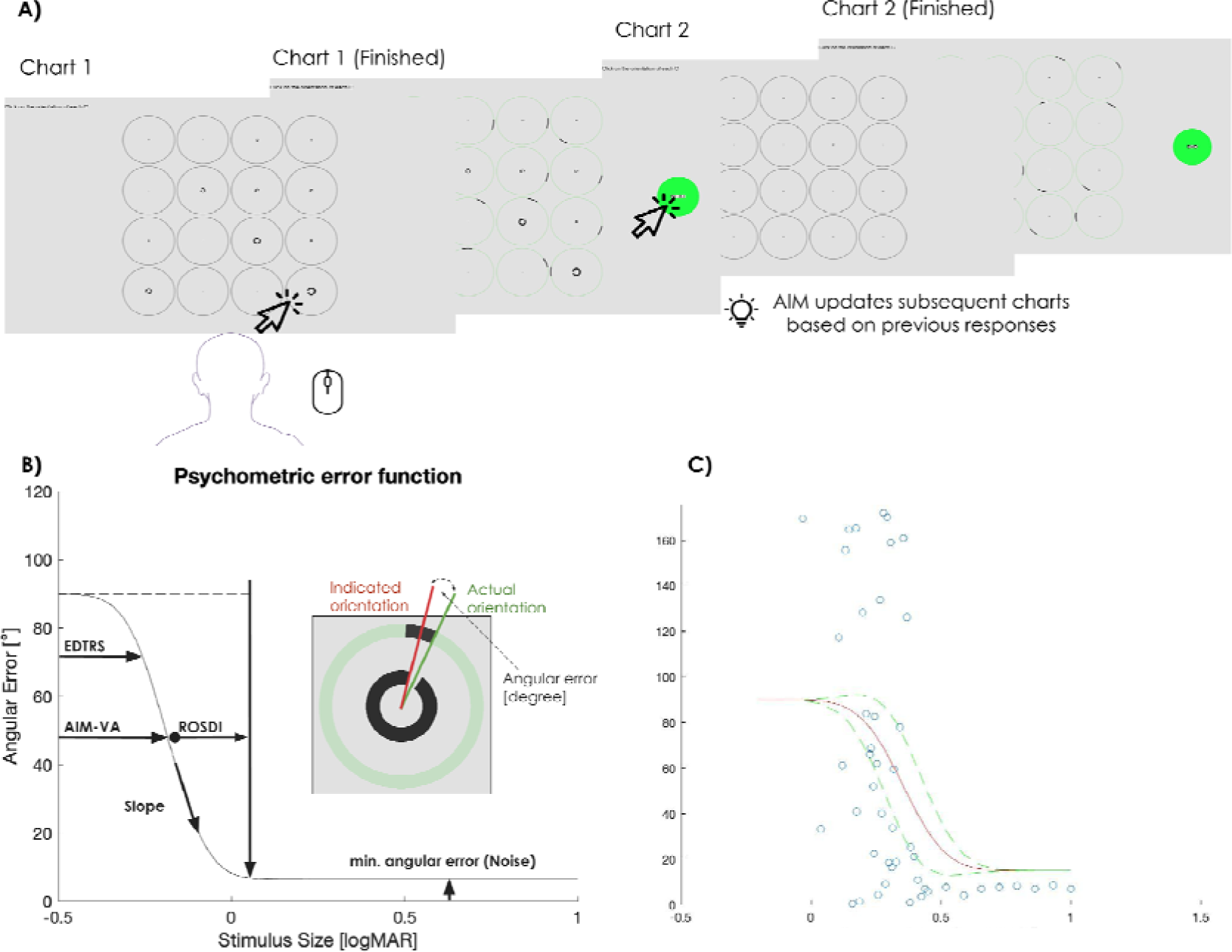
Schematic of AIM principal. A) In series of charts, each containing a 4*4 grid of cells, stimuli that range from easy to hard-to-detect are presented while being scaled logarithmically and randomized in regards of size and gap orientation. Each cell consists of a stimulus surrounded by a response ring. Participants reported the orientation of the C optotype in each cell by clicking on the corresponding location on the response ring. Their response is indicated by an arc on the ring, and participants are able to adjust their response by clicking again on the response ring. After a response has been made for all cells, a ‘Next’ button is presented, which participants can click when they are satisfied with their responses to the chart. The stimulus sizes on the next chart are updated, based on the responses to stimuli in previous charts and again logarithmically scaled in size. B) Schematic of AIM scoring and outcome parameters. Y axis represent the angular indication error, the x axis the stimulus size. C) Representative results and fit after three charts including psychometric function (red line), 95% confidence intervals (dashed green lines), and single responses (blue circles).

We presented a range of C optotypes with logarithmically spaced sizes. This feature, unlike single trial AFC methods, ensures that an observer always views stimuli that range from likely to unlikely to be seen and thereby ensuring for the inclusion of a clear exemplar, which may be particularly important for observers with visual or cognitive impairment. C optotypes were presented on a series of 2 charts, each comprising a 4×4 grid of 1.3° cells surrounded by a 0.1° thick indication ring. The background luminance and stimulus luminance were adjusted to match that of the ETDRS chart. The pixel size was 0.0026° (384 pixel/1°). The C gap width was kept at 20% of the line width and was constructed as an arc. The inter-cell gap distance was 0.1°. The surrounding rings had a grey hue and changed color to green to visualize an indication was made as well as the appearance of a black arc that indicated the reported gap orientation (Figure 1 A).

The stimulus sizes varied from difficult to easy, with log-step sizes starting from −0.30 logMAR to 0.80 logMAR on the first chart, following the design principle of the ETDRS chart (19). The range of sizes on the second and subsequent charts were in equal log steps between the latest threshold size estimate ±2σ from the fit to the results of previous charts. The orientation of the optotype in each cell was randomly drawn from a uniform distribution and the different optotype sizes were randomly assigned to each cell. Participants indicated the perceived orientation of all 16 stimuli on one chart by clicking on each ring (Figure 1 A), guessing if unsure. Participants could repeatedly correct their choice by clicking again on any response ring and proceeded to the next chart by clicking on a ‘Next’ button that became available only once responses had been made to all 16 stimuli. The response error is calculated as the difference between target orientation and indicated orientation in degrees (see Figure 1 B).

The response errors for each optotype size were converted to 8AFC responses, where absolute orientation errors less than 22.5° was scored as correct response, and errors greater than 22.5° were scored as incorrect responses. A cumulative gaussian psychometric function was fit to the 8AFC data, from which VA threshold ±2σ was used to select the optotype sizes for the next chart.

At the end of the final chart, a cumulative gaussian function was fit to the error report data as a function of optotype size from all charts, defined as:

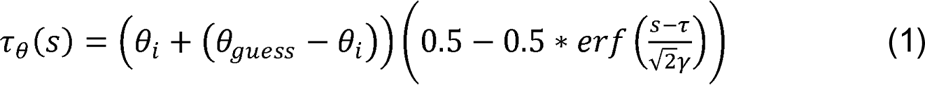

Where θ*_guess_* is orientation error for a guess response (90° for a stimulus with 360° range e.g., C; or 45° for a stimulus with 180° range e.g., grating-type stimuli), *s* is signal intensity (here stimulus size), τ is a sensitivity threshold, θ*_i_* is internal angular uncertainty (*noise* – the angular report error for the most visible stimulus), and γ is the slope of the function.

Figure 1 B) shows the different parameters in a schematic diagram of the psychometric fit after an AIM experiment. AIM’s threshold, here AIM-VA, is halfway between random responses and plateauing of performance, i.e., *noise*. As an existing test may have a fixed threshold criterion, e.g., ETDRS chart min. 60% correct, we can calculate an ETDRS-equivalent VA. To do so, we constrained the free parameters of AIM’s fit for noise and slope so that the number of free parameters was comparable to the ETDRS letter chart. On the other hand, AIM-VA can be additionally calculated with slope and noise as additional free parameters. This difference may cause slight variations in VA performance and also enable personalized models of psychometric fits (See Supplementary materials, Figure S1 for more details on fully versus semi-constrained fits). Specifically, the slope of a psychometric function reveals the change of the detection probability across stimulus levels, and is not scored for ETDRS although it can be used to report the logMAR range from guessing to perfect performance (29). AIM provides a similar range, which we define as the logMAR range between VA and the 95% point on the cumulative Gaussian function (5% above an observers min. angular indication error, Figure 1B). We refer to the logMAR distance between VA and the 95% angular error point as *range of stimulus detectability improvement* (ROSDI), which describes the range of optotype sizes over which visual function improves that may provide useful information concerning vision changes in different eye diseases. As threshold, slope, and noise interact, ROSDI is a single endpoint that incorporates all these parameters.

Figure 1C shows example data for one observer where the x axis shows optotype size in logMAR units, the y axis shows absolute angular error in degrees, blue circles show the orientation reports of an observer, the red line shows the best fitting cumulative gaussian function, green dashed lines show 95% confidence intervals at each optotype size.

Orientation errors resulting from an observer’s responses should be scattered randomly in either clockwise or anti-clockwise orientation relative to the target orientation and increase in magnitude with decreasing stimulus size. However, if optical distortion is present due to astigmatism, a systematic bias in error reports may occur that is expected to follow a sinewave pattern. If an observer is corrected for astigmatism, then distortions should be minimized unless neuro-ophthalmic distortion is present e.g., due to amblyopia (35). AIM can quantify orientation report error bias. The orientation bias was calculated as a function of error direction (clockwise or anti-clockwise relative to target orientation), target orientation and large to small stimulus sizes using a two-parameter sinewave function to calculate phase (bias) and its amplitude.

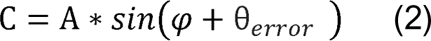

Where θ*_error_* is the observer’s orientation error for a target gap orientation, *φ* is the phase (bias), and *A* is amplitude of the function. Also, we calculated the R^2^ to evaluate the goodness of fit.

### 2.4 Procedure

To ensure comparable luminance levels, we measured at 1m the mean letter and background luminances for ETDRS with a LS 110 photometer, which were 5.2cd/m^2^ ±1.3 and 190.6 cd/m^2^ ±1.6, respectively. The minimum and maximum luminance values of the monitor were 0.3 cd/m^2^ and 250 cd/m^2^, respectively and gamma was 2.0. The LUT values for the background was adjusted to 191 to match AIM-VA to ETDRS.

#### Pilot experiment-refractive error methods

We fitted constrained psychometric functions (threshold as a free parameter for each fit, with slope and error shared across all fits for each observer) and calculated ETDRS-equivalent VA for AIM. The test was performed in 4m viewing distance at constant lighting conditions. Two charts, including one adaptive step, were deployed to measure the VA. One-D (Diopter)-step ±spherical lenses were placed in a trial-frame until the largest C was unrecognizable. We analyzed data ranging between −10.00D and +3.00D. Matlab’s *anova1* and *multcompare* (Tukey-Kramer test) functions were used for ANOVA analysis and planned comparisons, respectively.

#### First experiment

We used spherical (i.e., 0.00D, +0.25D, +0.50D, +0.75D, +1.00D, +2.00D) followed by cylindrical blur lenses (+0.50D, +1.00D, +2.00D, each in 0°, 90°, 135°) at a 4m viewing distance (thus +0.25D remaining accommodation). For each part of the experiment, lenses were randomized within and between observers and placed in front of the dominant eye in a conventional trial frame while the opposite eye was occluded with a dark occluding lens. Matlab’s *anova1*, *anovan,* and *multcompare* functions were used for ANOVA analysis and planned, multi-comparisons, respectively, for thresholds, noise, slope, ROSDI, and the phase and amplitude (we converted all values to positive numbers) values of the orientation report error analysis. All other aspects were the same as in the pilot experiment. For ETDRS, VA is defined at the line at which 3/5 (60% correct) letters are correctly identified, which corresponds to an angular error of ±72° for AIM-VA. ETDRS has a single parameter estimate (threshold -VA), to equate the number of free parameters we shared the slope and noise level parameters across all data for each subject and allowed threshold to vary in order to equate the number of free parameters for AIM-VA and ETDRS. The observer’s task during the ETDRS test was to read the letters until the line that was read with more than 2 incorrect answers. Each correct letter was count 0.02logMAR within the last line, e.g., if the performance at the last line was at 0.00logMAR and 4/5 correct answers were given, this would result in +0.02logMAR VA.

#### Second experiment

We investigated the repeatability of AIM-VA and compared it to ETDRS test-retest. The dominant eye was tested, and each test was repeated twice on the same day by the same experimenter where the order of tests was randomized using two charts for AIM.

#### Control experiment

To test whether more adaptive steps for AIM-VA would have an impact on repeatability, we performed a control experiment with a subset of participants who were tested with four adaptive steps (5 charts). Bland Altman plots were used in combination with a linear regression model.

## 3. Results

### 3.1 Experiment 1-Testing AIM’s ability to detect induced defocus and astigmatic blur

We initially performed a pilot experiment with negative and positive spherical 1.00D lenses in young (mean age 21), normally sighted observers. A one-way ANOVA (−10.00D to +3.00D) revealed a significant effect of defocus on VA [F(13,140)=14.0, p<0.001, η_p_^2^= 0.57]. All positive spherical lenses caused a defocus-based decline in VA with a significant effect using +2.00D (planned comparisons). Negative lenses caused an accommodation-based flat function of VA until approximately −5.00D, and then VA decreased significantly for-10.00D. No significant effect of defocus on psychometric function slopes [F(13,140)=0.89, p=0.57, η_p_^2^= 0.08] and ROSDI [F(13,140)=0.89, p=0.57, η_p_^2^= 0.08] was found but there was a significant increase in orientation error noise [F(13,140)=3.86, p<0.001, η_p_^2^= 0.26] due to induced refractive error. A planned comparison showed this was due to positive blur (see supplementary materials, Figure S2 & S3).

We then investigated whether AIM would be able to detect defocus blur smaller than 1 diopter and compared results with that of an ETDRS letter chart.

For the group without correction worn, the median spherical (sph), negative cylindrical (cyl) values in diopters, axis (x) in degrees, spherical equivalent (SE) in diopters, and their respective minimum to maximum values were:

OD-sph: +0.13 D [-0.25, +1.00] / cyl: −0.75 D [-0.75, ±0.00] x 94° [89, 164] | SE: −0.19 D [-0.75, +0.63]

OS-sph: ±0.00 D [-0.25, +1.75] / cyl: −0.38 D [-1.25, ±0.00] x 65° [9, 179] | SE: ±0.00 D [-0.75, +0.63].

The effect of spherical blur induced by using positive spherical lenses is shown in Figure 2. A two-way ANOVA between test types and blur levels revealed a significant effect of test type [F(1,252)=24.0,p<0.001, η_p_^2^= 0.09] and spherical blur [F(5,252)=43.7,p<0.001, η_p_^2^= 0.46], but no interaction between these independent variables [F(5,252)=1.4,p>0.05, η_p_^2^= 0.03]. Planned multiple comparisons between the +0.00D and all other blur conditions within each test type showed an elevated, i.e., poorer, but non-significant acuity for +0.25D and +0.50D and a significant reduction in acuity with +0.75D but with a stronger effect for AIM [p<0.001] compared to ETDRS [p<0.05] and +1.75D blur for both AIM and ETDRS [p<0.001].

**Figure 2:**
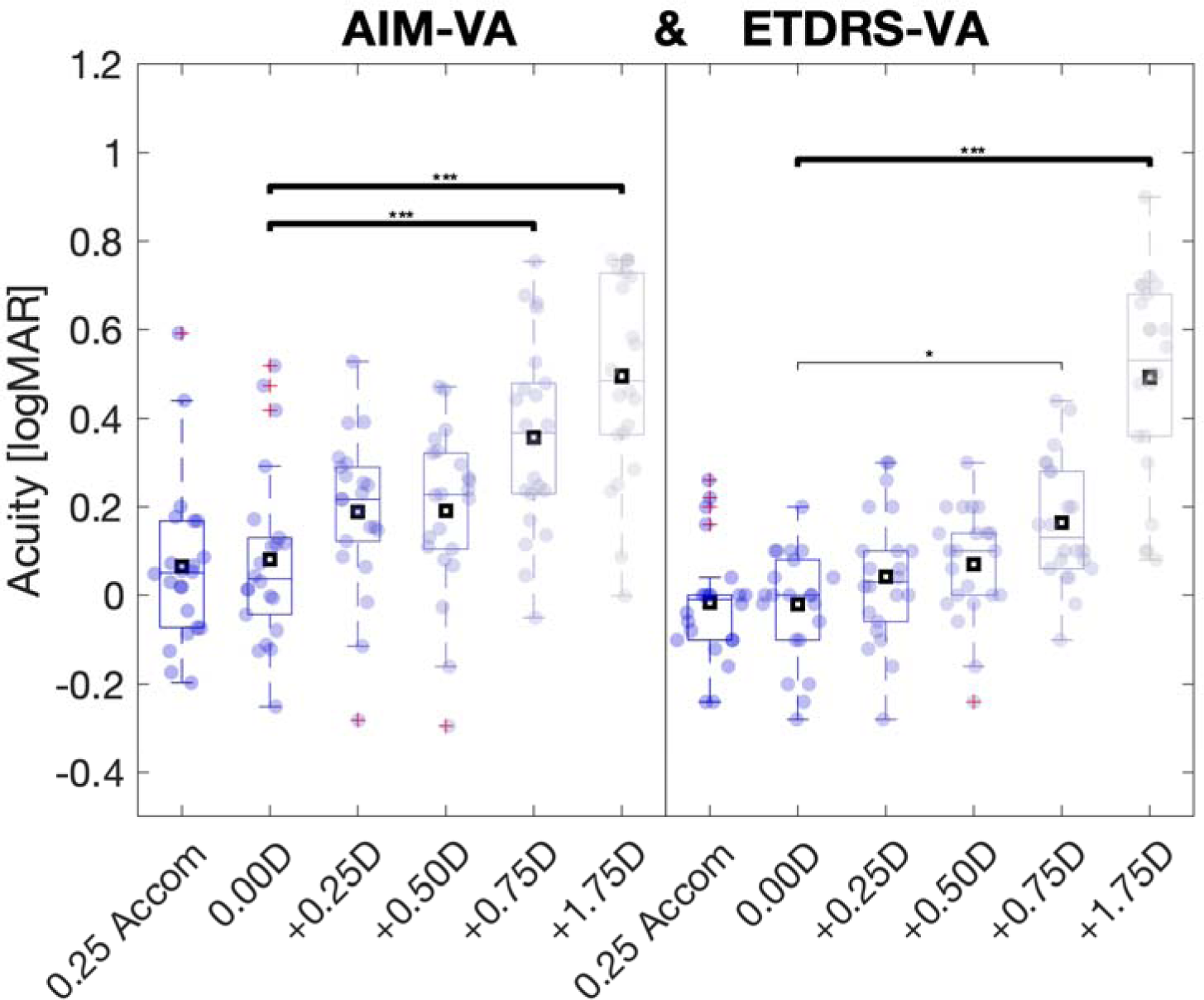
Results for induced spherical defocus results for AIM (left) and ETDRS (right). Shown are interquartile range (boxes), medians and means (horizonal lines and squares within each box, respectively), single outcomes (dots) jittered by a kernel density estimate horizontally, whiskers for extreme values, and red + for outliers. Depicted are VA results (y axis) for each effective blur level at 4m distance (i.e., lens power minus 0.25D accommodation), using plano lens, +0.25D, +0.50D, +0.75D, +1.00D, +2.00D spherical lenses. *Indicated p<0.05; *** p<0.001 of planned comparisons between 0.00D and blur conditions.

We also tested pairwise differences between test types for each blur condition using planned multiple comparison analysis and found that all AIM VA means were nominal higher but only the +1.00D lens (effective +0.75D blur) was significantly different [p<0.05]. The mean differences (AIM-ETDRS) were all positive (supplementary materials, Table 1).

**Table 1.** Comparison of absolute and relative (%) test-retest differences across levels of difference in logMAR as a function of number of adaptive steps and number of charts during the control experiment. Bias is the mean of the difference between test and retest results.

| AIM-VA<br>Chart/<br>Adaptive<br>steps | Bias | $\leq \pm 0.10$<br>logMAR<br>[% \| n] | $\leq \pm 0.20$<br>logMAR<br>[% \| n] | $\leq \pm 0.30$<br>logMAR<br>[% \| n] | $\leq \pm 0.40$<br>logMAR<br>[% \| n] | Limits of<br>agreement<br>[logMAR] |
| --- | --- | --- | --- | --- | --- | --- |
| 1/0 | 0.039 | 25% \| 3/12 | 58% \| 7/12 | 92% \| 11/12 | 92% \| 11/12 | 0.44 / -0.37 |
| 2/1 | 0.007 | 42% \| 5/12 | 92% \| 11/12 | 100% \| 12/12 | 100% \| 12/12 | 0.30 / -0.28 |
| 3/2 | 0.002 | 50% \| 6/12 | 92% \| 11/12 | 100% \| 12/12 | 100% \| 12/12 | 0.24 / -0.23 |
| 4/3 | 0.015 | 75% \| 9/12 | 100% \| 12/12 | 100% \| 12/12 | 100% \| 12/12 | 0.18 / -0.15 |
| 5/4 | 0.023 | 75% \| 9/12 | 100% \| 12/12 | 100% \| 12/12 | 100% \| 12/12 | 0.16 / -0.12 |

We also investigated whether spherical blur affects the other parameters and found that the minimum orientation error (noise) was significantly increased [F(4.0)=131,p<0.01, η_p_^2^= 0.12] with blur whereas the slope was unchanged [F(0.4)=131,p>0.05, η_p_^2^= 0.01]. ROSDI (see Figure 1B) includes the distance between 5% above an individual’s minimum angular error (noise) to the threshold and depends also on the slope of the psychometric function. Therefore, it contains all three parameters, which may interact with each other. We investigated the effect of spherical blur and found no significant change of ROSDI [F(5,131)=0.4, p>0.05, η_p_^2^= 0.01], (supplementary materials, Figure S4, S5, S6).

The median duration for AIM across blur conditions using two charts was 59 ±25 (standard deviation) sec. We did not collect data for time across blur steps for ETDRS and found for AIM no significant effect of blur on experimental duration [F(1,131)=0.9, p>0.05, η_p_^2^= 0.03] (supplementary materials, Figure S7).

Next, we investigated the effect of astigmatic blur. A 3-way ANOVA was performed and there was a significant effect of test type [F(1,382)=63.7,p<0.001, η_p_^2^= 0.14], blur level [F(2,382)=72.0,p<0.001, η_p_^2^= 0.27], and cylindrical axis orientation [F(2,382)=8.4,p<0.001, η_p_^2^= 0.04] but no interaction effects. Planned multiple comparison analysis between test types revealed that many AIM VAs, including all 90° conditions, were significantly lower (numerically higher) compared ETDRS and all mean differences (AIM-EDTRS) were positive (supplementary material, Table 2).

A two-way ANOVA showed no significant effect of blur level [F(2,189) =1.9,p>0.05, η^p2^= 0.02], cylindrical axis orientation [F(2,189)=0.7,p>0.05, η_p_^2^= 0.007] nor an interaction [F(4,189)=0.3,p>0.05, η_p_^2^= 0.006]. (supplementary materials, Figure S8). Noise showed a significant effect of blur level [F(2,189)=11.3,p<0.001, η_p_^2^= 0.10] and cylindrical axis orientation [F(2,189)=4.1,p<0.05, η_p_^2^= 0.04], but no interaction [F(4,189)=0.6,p>0.05, η_p_^2^= 0.02](supplementary materials, Figure S9). No significant effects of ROSDI for cylindrical blur condition [F(2,189)=1.9, p>0.05, η_p_^2^= 0.02], axis orientation [F(2,189)=0.7, p>0.05, η_p_^2^= 0.007], or interaction between both [F(4,189)=0.2, p>0.05, η_p_^2^= 0.006] were found (supplementary materials, Figure S10).

The median duration for administering AIM using two charts and using cylindrical blur was 46.8 sec ±17.7 (supplementary materials, Figure S11).

### 3.2 Analysis of orientation report error

To confirm that the model is insensitive to defocus blur, we analyzed whether spherical lens induced blur would affect the two parameters of the sinewave model using one-way ANOVAs and found that neither phase [F(5,114)=0.7,p>0.05, η_p_^2^= 0.03] nor amplitude [F(5,114)=1.8,p>0.05, η_p_^2^= 0.07] were significantly affected (supplementary materials, Figures S12, S13).

We then tested whether cylindrical lenses would systematically change the orientation report error, using a sinewave model that accounts for optical distortion of perceptual to actual location of the gap (Example Figure 4). First, we analyzed the phase shift using two-way ANOVAs. We found no significant effect of blur [F(2,171)=0.2,p>0.05, η_p_^2^= 0.003], no effect of axis orientation [F(2,171)=0.5,p>0.05, η_p_^2^= 0.006], nor an interaction [F(4,171)=0.4, p>0.05, η_p_^2^= 0.009] for the phase parameter. The amplitude of the model was significantly affected by cylindrical blur [F(2,171)=9.2,p<0.001, η_p_^2^= 0.08] i.e. higher blur caused larger amplitude, but not by axis orientation [F(2,171)=0.9,p>0.05, η_p_^2^= 0.006], nor was an interaction found [F(4,171)=1.4,p>0.05, η_p_^2^= 0.02] (supplementary materials, Figures S14, S15).

### 3.3 Experiment 2-Repeatability of AIM-VA

The Bland-Altman plots in Figure 5 show repeatability for AIM-VA using one adaptive step (2 charts; 5A) and ETDRS (5E) during the second experiment and the results for the control experiment (5B-D) with two to four adaptive steps. Test-retest VA differences (y axis) were larger for AIM-VA than ETDRS using one adaptive step, but not for two or more adaptive steps. AIM’s mean VA test-retest bias was −0.03 using one adaptive step and +0.04 for ETDRS. The relative and absolute numbers of test-retest difference for AIM-VA were ≤0.10logMAR were 68% (15/22) and ≤0.20logMAR 86% (19/22) and for ETDRS ≤0.10logMAR were 86% (19/22) and ≤0.20logMAR 100% (22/22). Limits of agreement in logMAR for test-retest difference were AIM-VA [-0.36 0.41] and for ETDRS [-0.11 0.18].

As the variance of an AIM fit depends on the amount of data, its variance should decrease with increasing trials. We ran a control experiment with four adaptive steps (5 charts in total in a subset of twelve participants to test this hypothesis. As shown in Figure 5B-D, the distance between upper and lower limits of agreement in the Bland Altman plots significantly decreased with step sizes [t(4) =5.2, p<0.01] and were comparable to that of ETDRS. We calculated whether and if so, how much the test-retest difference improves across trials in absolute and relative numbers as well as by determining the upper and lower limits of agreement (Table 1).

## 4. Discussion

### 4.1 AIM’s ability to detect induced defocus and astigmatism

One of the objectives of the current study was to test AIM-VA’s ability to detect refractive errors and compare that ability with a standard letter chart. We show that AIM detects defocus blur and that it is as sensitive to detect blur as the ETDRS chart (Figure 2). A previous study tested the effect of spherical and astigmatic blur induced with lenses on E and C using +1.50D (36), which is comparable with the loss of vision shown in our study. We also show that oblique induced astigmatism has a stronger blur effect than with or against the rule astigmatism (Figure 3), in good agreement with a previous study (37). Interestingly, when comparing AIM and ETDRS in regards of detecting astigmatic blur, AIM showed larger loss of VA. Different optotypes vary in their spatial frequency contents (38, 39). letters other than uniform stimuli like the C or O varying more with respect to orientation-specific spatial frequency content. The combination of the continuous error report and the more homogenous spatial frequency properties of the C used for AIM maybe more suitable to detect astigmatic blur compared to optotypes in the EDTRS chart that are more heterogenous regarding orientation and spatial frequency properties. AIM showed overall more intra-individual variability for both defocus and astigmatism than ETDRS, which may be due to the larger degrees of freedom due the continuous indication task and 360-degree randomization of the C gaps compared to the five-letters-per-line letter identification task.

**Figure 3:**
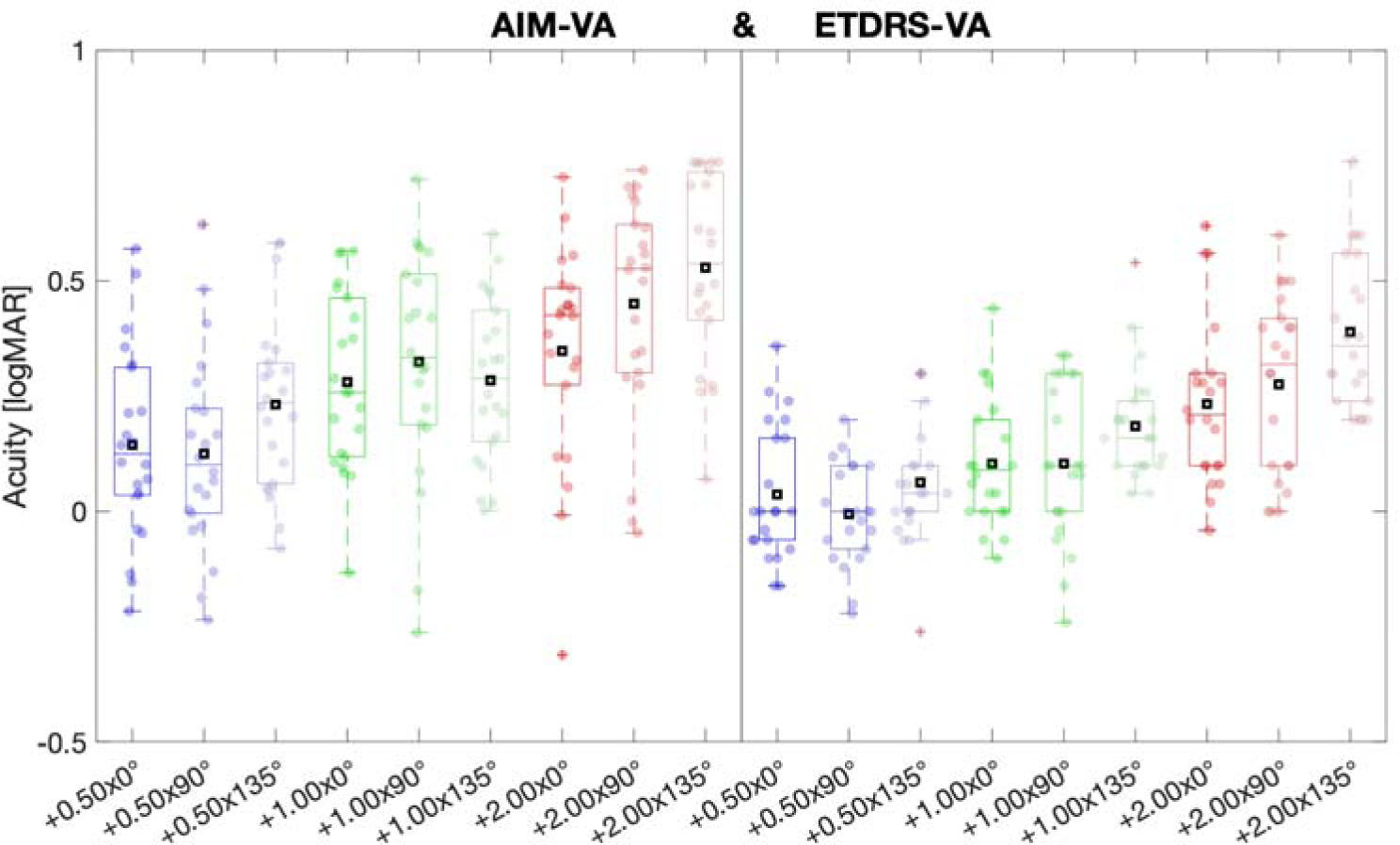
Results for AIM (left) and ETDRS (right) VAs using cylindrical lenses. Depicted are VA results (y axis) for each blur lens (x-axis), i.e., +0.50D (blue), +1.00D (green), and +2.00D (red) cylindrical powers in combination with 0°, 90°, 135° induced orientations.

Clinical standard chart tests deliver threshold values that are known as VAs. Psychophysically, VA is one point on a psychometric function that describes the probability that an observer can detect a signal, in this study gap orientation or letter identity. The stimulus range over which an observer’s performance transitions from detecting to not being able to detect the signal is expressed by the slope of the psychometric function and may vary between observers and may provide additional important information about the health of the participant, the progression of eye disease or their response to treatment. Practically speaking, the slope describes the rate of transition from seeing to not seeing and can be either very steep or shallow. Chart tests, such as the ETDRS and Snellen charts, do not take this parameter into consideration whereas AIM-VA calculates a personalized slope value and generates a personalized VA using a psychometric function fitting approach (Figure 1C).

Moreover, even though two observers may be able to accurately report a signal in a forced choice paradigm, the quality of their vision may still be distinct between observers or across time. For example, in terms of a conventional letter chart, one observer may correctly name all five letters in all lines before not being able to detect any single letter whereas a second observer may reach the same VA but make mistakes in some of the lines with larger stimuli. Clinically those VAs are identical but the suprathreshold performance differed. Unlike letter charts such as ETDRS, AIM-VA provides an output for that observer-specific behavior, namely the minimum angular error parameter for the most intense (here: largest) stimuli. A number of sources can affect this parameter e.g., hand tremor, eye movements, or neurological intrinsic variations (spiking variations, neuro-transmitter variation etc.) and structural irregularities in the optics or retina. We termed these sources as *noise* that captures all sources of those external and internal variability. We show that this noise parameter significantly increased with increasing defocus and astigmatic blur.

Previous research demonstrated that the slope of the psychometric function is affected by blur (29), specifically that the slope becomes more shallow with one diopter defocus blur. Since threshold, slope, and noise interact, it remains unclear what causes a change in an observer’s psychophysical performance. Hence, we introduce the *range of stimulus detectability improvement*, ROSDI (Figure 1B), which takes all three parameters into account, ranging from the midpoint between random performance and plateauing visual performance (noise) to 5% above an individual’s noise value. ROSDI is expressed here in logMAR and can be converted to all other known VA units as well as across other signal intensity parameters. We found no systematic effect for ROSDI due to induced refractive error in this sample of healthy young observers, however, future research may consider ROSDI in pathological eye diseases.

Astigmatism is a prevalent condition worldwide, ranging in prevalence between 8 to 62% (40) and may induce perceptual distortions that affect performance during an acuity tasks. It has been shown that induced astigmatism generates neural response biases (41) and that observers may show intrinsic bias preferences (41, 42). We found that the induced error using astigmatic blur caused a significant effect of amplitude, i.e., increasing blur increased the amplitude, on the sinusoidal pattern of response errors, suggesting that the report error was systematically affected by astigmatism (Figure 4, see also Supplementary Materials, Figure S15). Future studies may use responses in a separate domain (e.g., verbal or manual) to estimate the relationship between report error and amplitude and phase under astigmatism.

**Figure 4:**
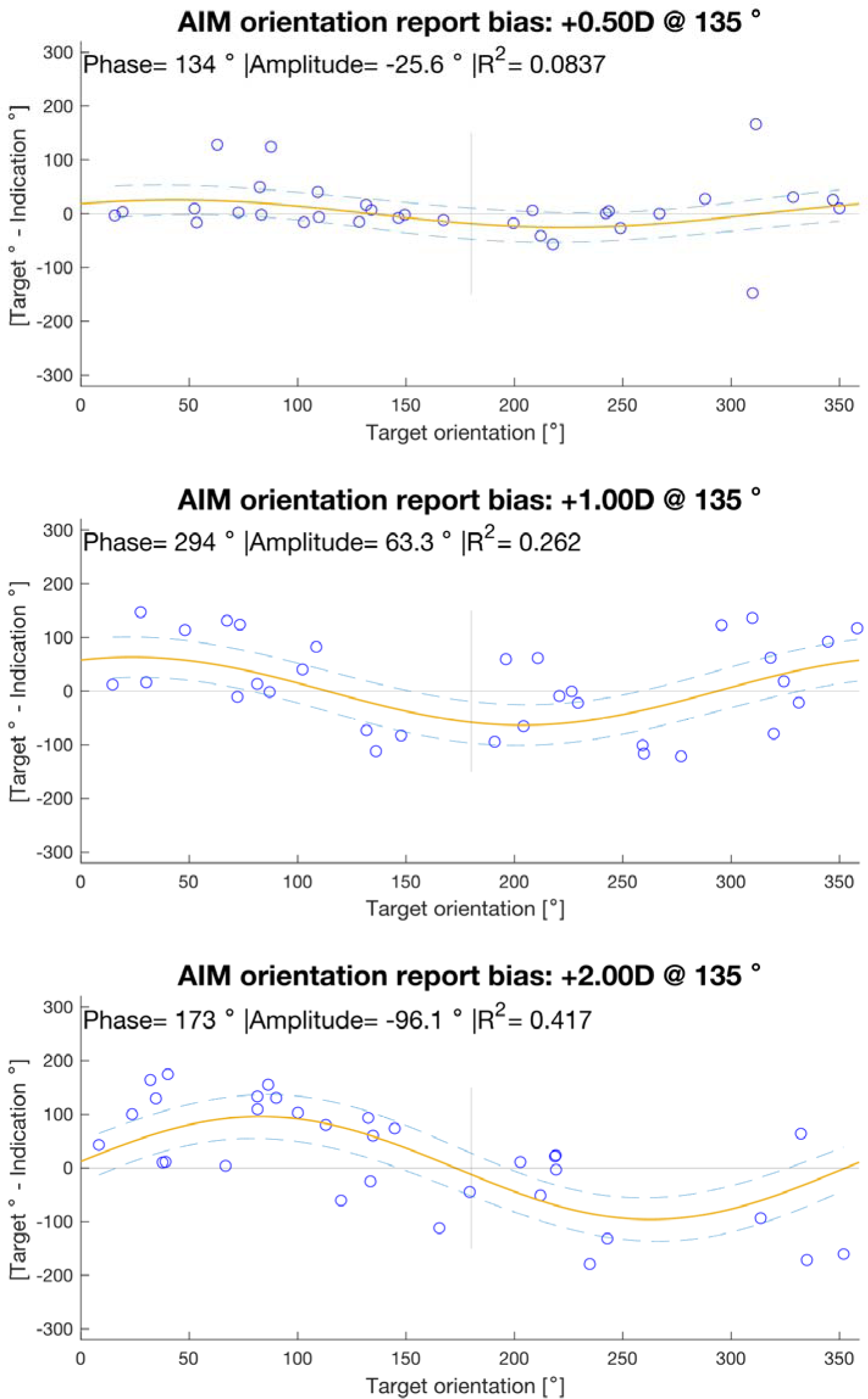
Example for an individual’s results applying the oblique cylindrical +0.50 D (top), +1.00D (middle), and +2.00D (bottom) blur. The orientation of each target ring was randomized. The y axis shows orientation report error, the x axis shows the target orientation. The orange line represents the best fitting sinewave function, the blue dashed lines show the 95% confidence intervals, single indications are shown as blue circles. The phase, amplitude, and goodness-of-fit R^2^ are depicted above.

### 4.2 AIM test-retest results

In experiment two, we compared AIM-VA with ETDRS using a single chart as often deployed in clinical practice. We show that AIM-VA using two charts (i.e., the initial chart and one adaptive step) has greater test-retest variance (Figure 5A and 5E). This may be due to three reasons, namely a memory bias for a single ETDRS chart, fewer decision options for ETDRS (10 letters) compared with AIM (continuous 360°), and AIM-VA’s sampling of data along the slope from random report error (90°) to the min. angular error. To address the latter point, we carried out a control experiment whereby we increased the number of adaptive steps to sample more data along the psychometric function to improve the fit. We hypothesized that within the time window of the test to retest (within one day), the slope, noise, and threshold of an observer should not significantly change. Indeed, increasing the number of charts improved the repeatability of AIM-VA significantly and was comparable with ETDRS using three or more adaptive steps charts with 4*4 cells (Figure 5B-D). The repeatability of ETDRS has been reported and 87% of the standard ETDRS chart generated VAs had a ≤0.10logMAR difference and 100% with a ≤0.30logMAR difference (43). Other tests such as FrACT reported comparable limits of agreement of approximately 0.20logMAR with a few outliers (44). AIM-VA showed comparable results when using two or more adaptive steps (Table 1).

**Figure 5:**
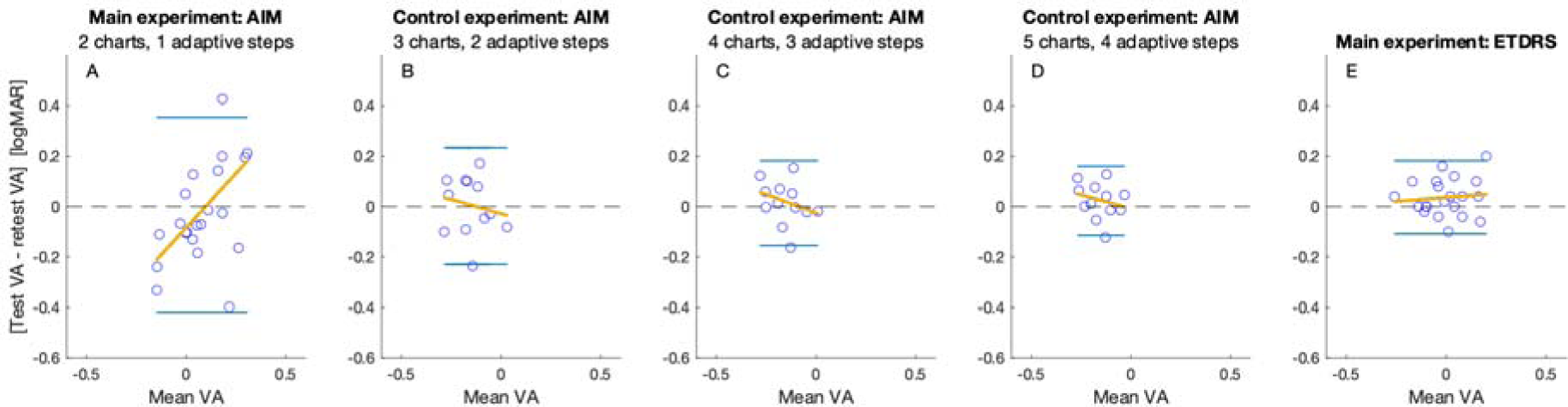
Bland-Altman plots for repeatability comparing AIM-VA with an initial chart and one adaptive step (A) and ETDRS with the same single chart repeated once (E) from the second experiment. Also shown are the control experiment results with two (B), three (C), and four (D) adaptive steps. Each y axis shows the VA test-retest difference in logMAR, each x-axis depicts the mean VA for test and retest VA results. Single circles represent data from individual participants, blue horizontal lines represent limits of agreement for test-retest VA difference, orange lines represent the linear function fits.

### 4.3 Future directions and study limitations

We acknowledge a number of limitations of the current study. We measured induced refractive error in a relatively small cohort to proof-of-concept. Future studies may explore the feasibility in a larger cohort, with and without visual impairments. In an ongoing study concerning VA in amblyopic children, preliminary results suggest that a near version of AIM-VA may be suitable to investigate vision with reliability (45). The positions of AIM-VA’s log-spaced stimuli sizes were randomized within each chart whereas ETDRS used log-spaced stimuli sizes that were organized from large to small down the chart. Future studies will examine whether an observer’s performance is significantly influenced by the randomization of the positions of letter sizes within each chart. We introduced a number of AIM-specific outcome measures including noise and ROSDI and show that noise was significantly increased with blur levels whereas no effect for ROSDI was found. Building a personalized model and using ROSDI longitudinal within observers is a future direction to investigate whether only single aspects, i.e., threshold, slope, or min. error or the entire model of the detectable range of visible stimuli may be affected, e.g., during amblyopia treatment. With AIM, the number of cells per chart and the number of adaptive steps can be independently varied. It is unclear how the repeatability changes by lowering the number of cells-per-charts while increasing the number of adaptive steps. Future research will investigate this question. In the current study, we applied AIM to measure VA, however, due to its generalizability, future studies will exchange stimulus size with other stimulus properties that are processed in distinct neural sites, e.g., color, contrast, stereo disparity, motion etc.

## 5. Conclusions

The Angular Indication Measurement, or AIM, is a response-adaptive, self-administered, and generalizable technique to interrogate visual functions. The current study applied AIM to measure the visual acuities in healthy adults and compared it with a standard clinical test in terms of its ability to detect induced refractive errors. We show that AIM showed comparable sensitivity to a conventional ETDRS letter chart. Using one adaptive step, AIM showed greater test-retest variance than a single ETDRS chart, which is likely due to memory bias and fewer degrees of variability for the ETDRS (10 letters) chart compared to AIM (continuous 360°). As hypothesized, increasing the number of adaptive steps improves the repeatability to the level of a single ETDRS chart. Based on the findings of the current study, at least 2 adaptive steps should be used to measure distance VA in adults reliably. An AIM chart took approximately 30sec to administer. As it provides additional information about an observer’s principal error level (noise), error bias, slope that considers an observer’s personalized psychometric function, and the range of stimulus detectability improvement, or ROSDI, in clinically useful VA units, e.g., logMAR, AIM may be a useful tool for researchers and clinicians to measure visual acuity.

## Acknowledgment

Supported by NIH R01 EY029713. Portions of this study have been presented at the virtual Vision Science Society Conference 2022 and Vision Science Society conference 2023.

## Data availability

Data underlying the results presented in this paper are not publicly available at this time but may be obtained from the authors upon reasonable request.

