## Supplementary materials for "AIM (Angular Indication Measurement)- Visual Acuity: An adaptive, self-administered, and generalizable vision assessment method used to measure visual acuity"

*Corresponding Author **

**Semi- versus fully-constraint psychometric fits**

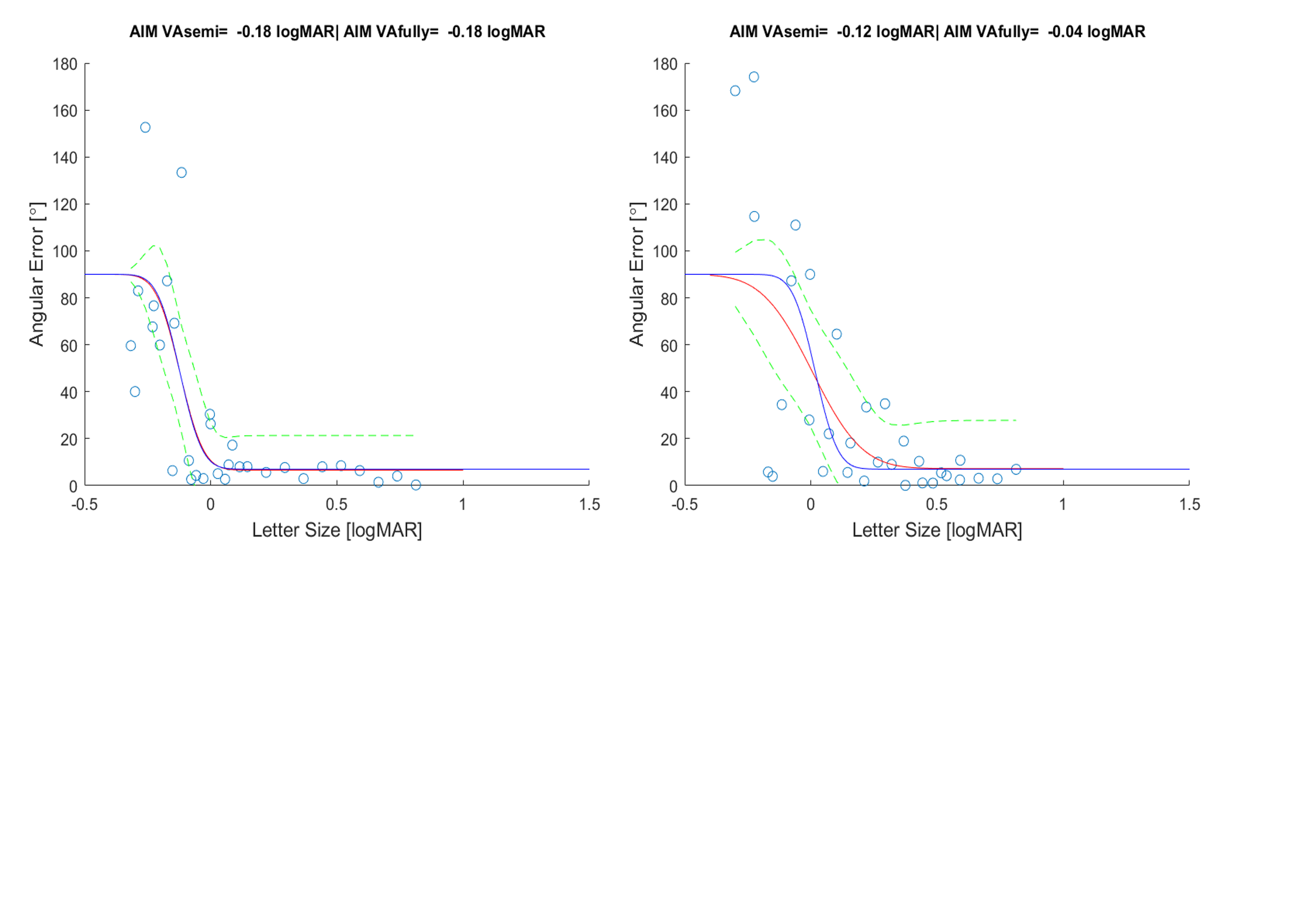

*Figure S1: Impact of the number of free parameters on psychometric function fits. Standard Acuity estimates (e.g. ETDRS) return a single parameter estimate, whereas AIM results are based on the fits of a psychometric function with 3 free parameters (equation 1in the main manuscript), which increases variance on the estimate of each parameter. To decrease the difference between methods, we compared a full model with 3 free parameters per psychometric function (right panel) with a constrained model in which the slope and minimum report error parameters were shared across all the data for a single observer (left panel).*

### Pilot experiment

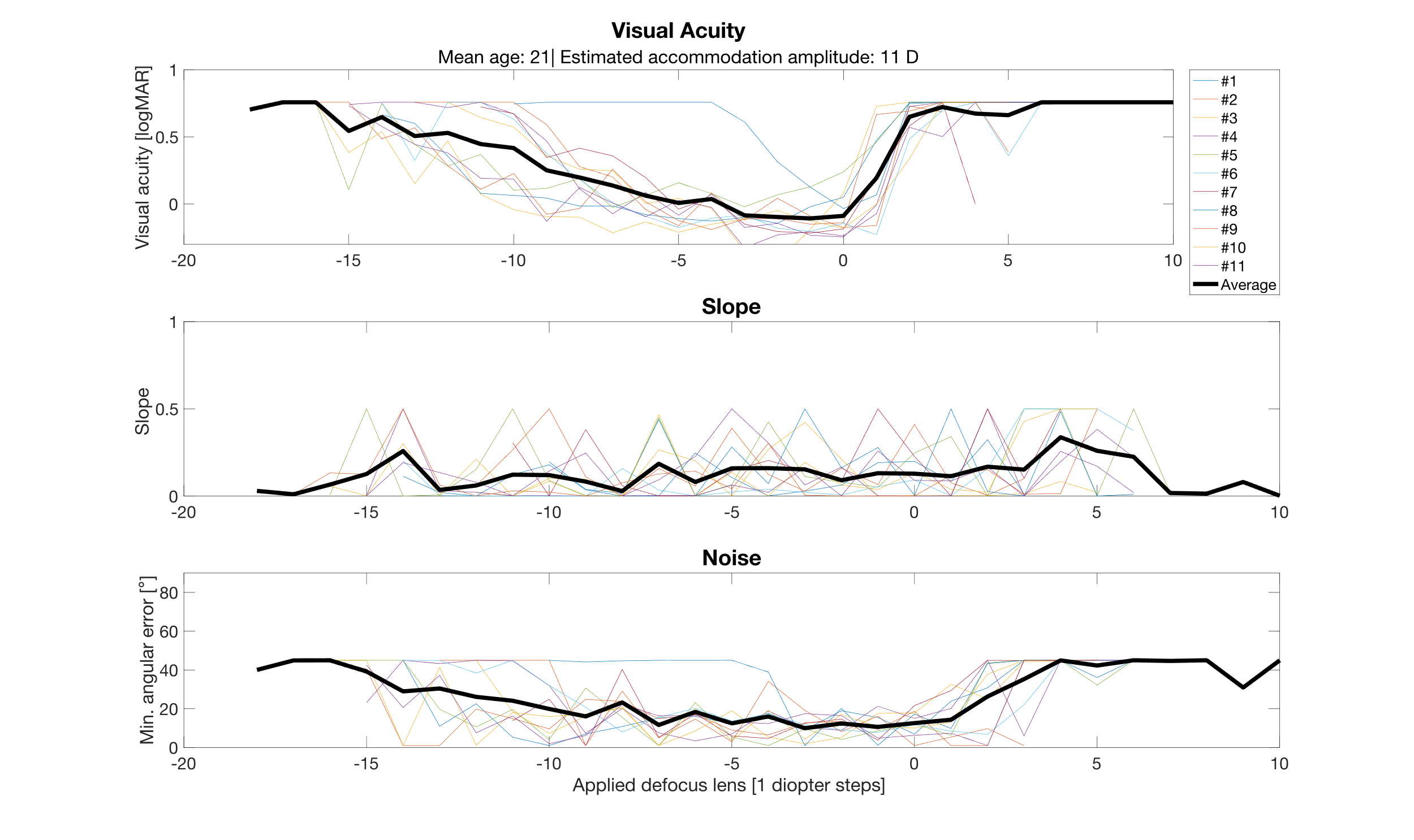

*Figure S2:* AIM VA psychometric function parameters (graph subtitle) as a function of defocus from the pilot experiment. X axis shows the added defocus in diopters, y axis shows the parameter values for VA (top), slope (middle), & noise (bottom). Data for individual observers are shown as colored lines, the group mean is shown by the black lines.

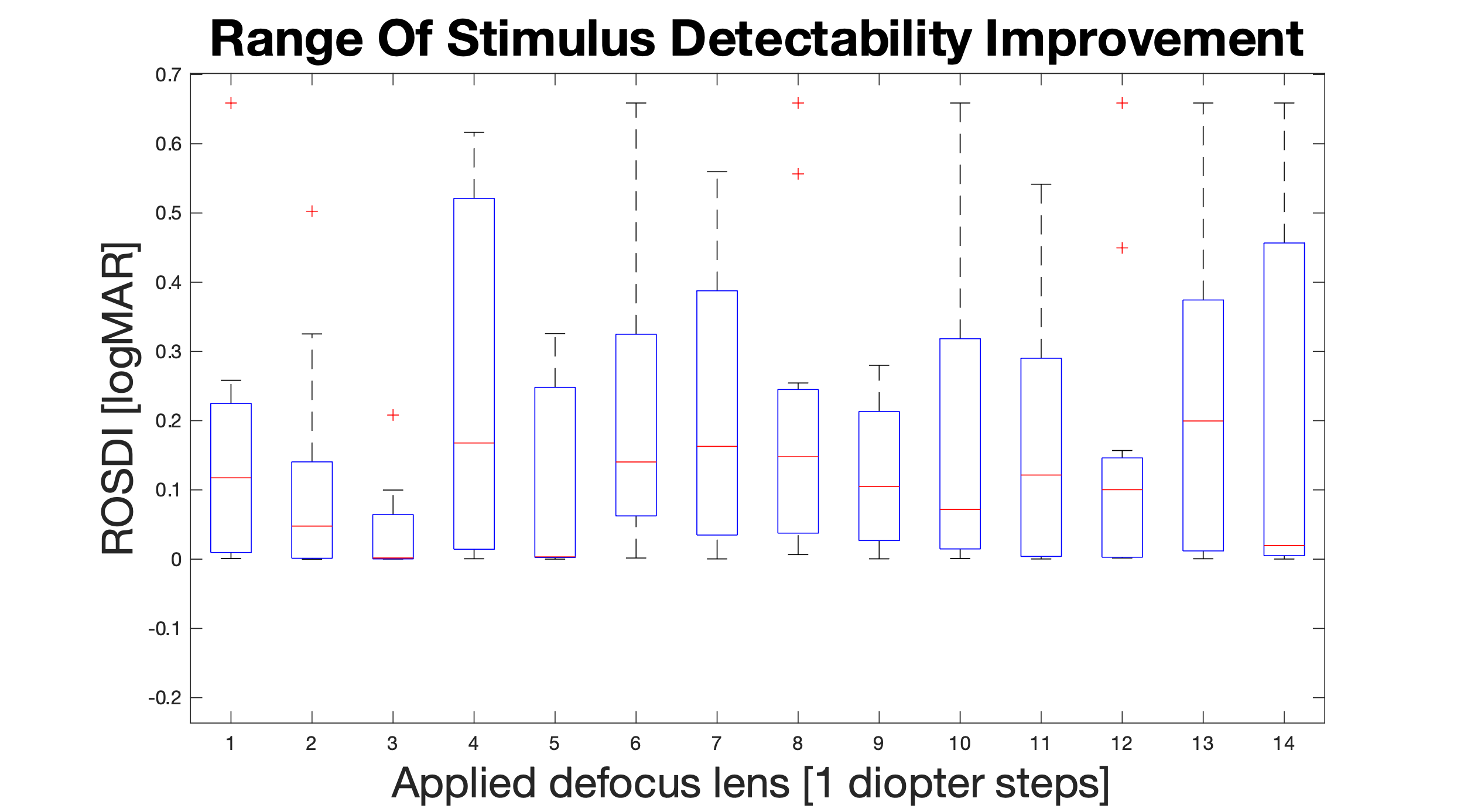

*Figure S3: Range of stimulus detectability improvement (ROSDI) as a function of blur for the pilot experiment, details as for figure S2, except for the ROSDI estimate from the psychometric function.*

### Experiment 1: Testing AIM’s ability to detect induced defocus

***Table S1: Planned multiple comparison results for difference between AIM and ETDRS under induced defocus using spherical lenses. Shown are comparisons between AIM and ETDRS values for each blur condition, including 95% confidence intervals, differences of means between each stimulus type, and p-values for testing for differences.***

| **AIM vs. ETDRS [D]** | **Lower 95%CI** | **Difference means** | **Upper 95%CI** | **P-value** |
| --- | --- | --- | --- | --- |
| **0.00** | -0.0938019747409674 | 0.0823081105618857 | 0.258418195864739 | 0.933326286320034 |
| **+0.25** | -0.0743221578015899 | 0.101787927501263 | 0.277898012804117 | 0.766539467279397 |
| **+0.50** | -0.0291486613058074 | 0.146961423997046 | 0.323071509299899 | 0.212731828288130 |
| **+0.75** | -0.0549077902938554 | 0.121202295008998 | 0.297312380311851 | 0.513508817035103 |
| **+1.00** | 0.0165111377038837 | 0.192621223006737 | 0.368731308309590 | **0.0182243855268087** |
| **+2.00** | -0.173488714373390 | 0.00262137092946274 | 0.178731456232316 | 0.999999775611513 |

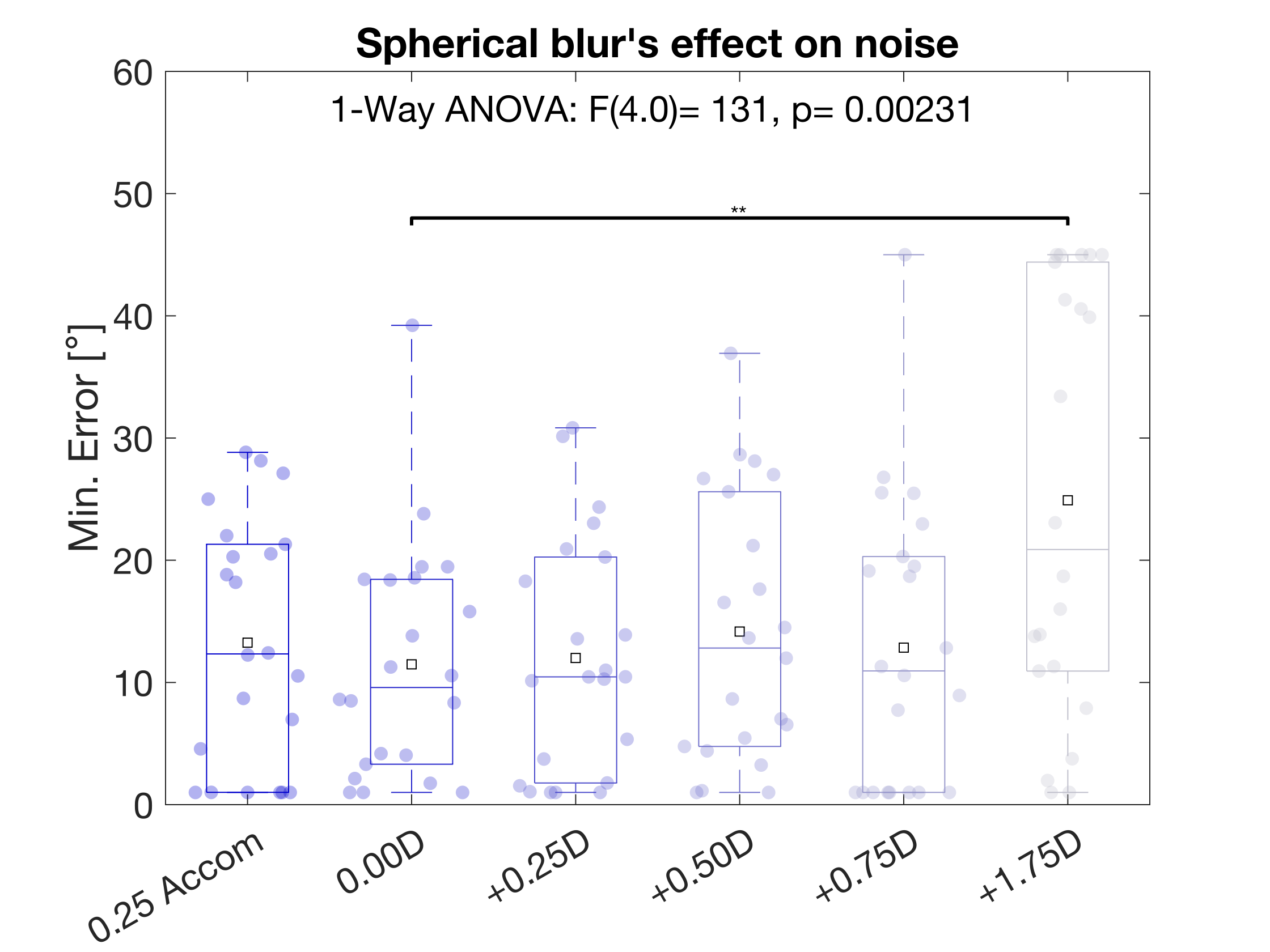

*Figure S4: Effect of spherical blur on minimum orientation report error of the data show min. angular error as a function of spherical blur levels in Experiment 1. Shown are interquartile range (boxes), medians and means (horizonal lines and squares within each box), single outcomes (dots) jittered horizontally by a kernel density estimate, whiskers indicate 95% confidence intervals, and + symbols show outliers. A one-way ANOVA showed a significant effect of spherical blur, and a planned multi-comparison revealed a significant difference between 0.00D and +1.75D condition.*

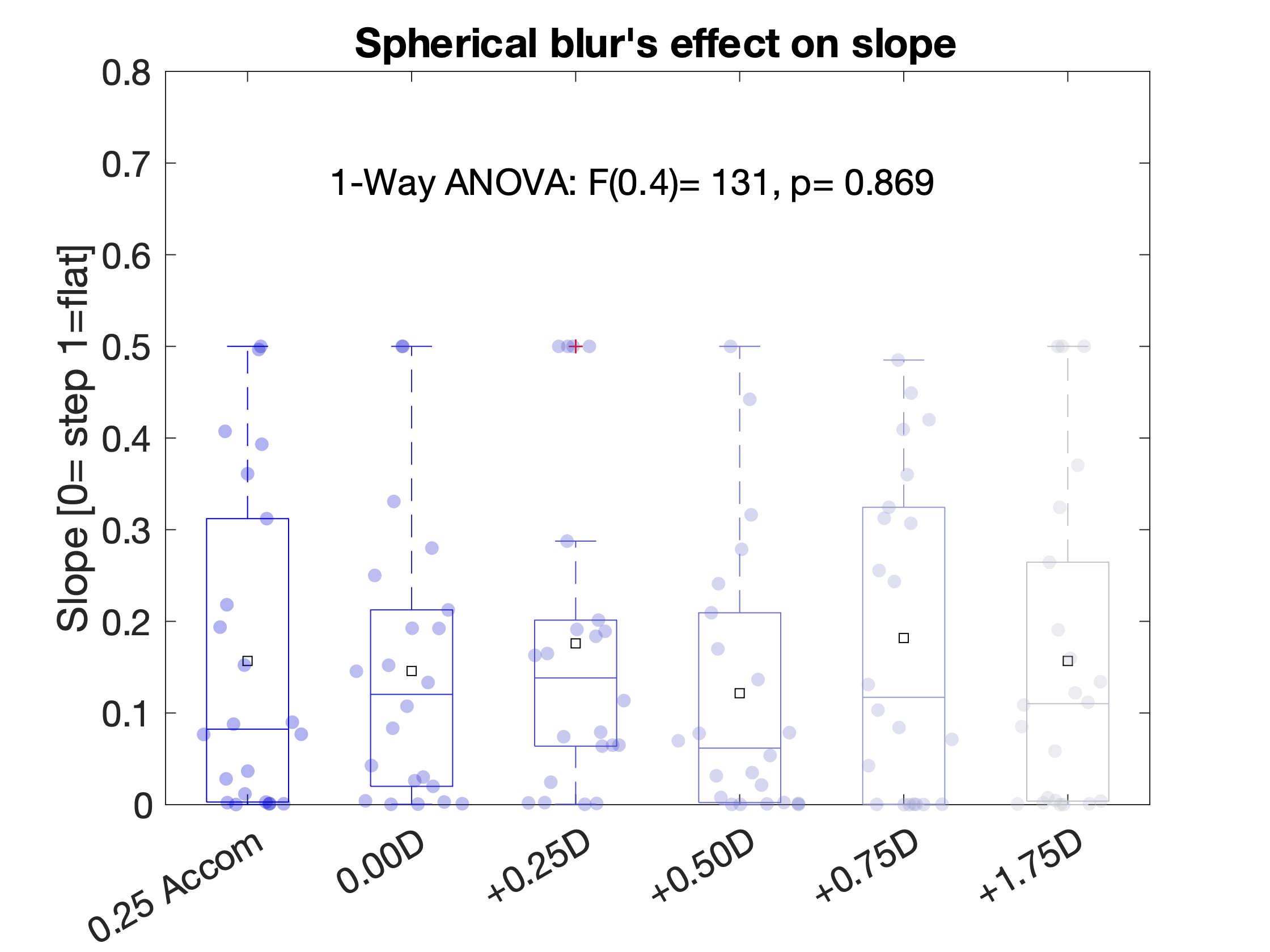

*Figure S5: Effect of Spherical Blur on the slope of the psychometric function. A one-way ANOVA showed no significant effect of spherical blur.*

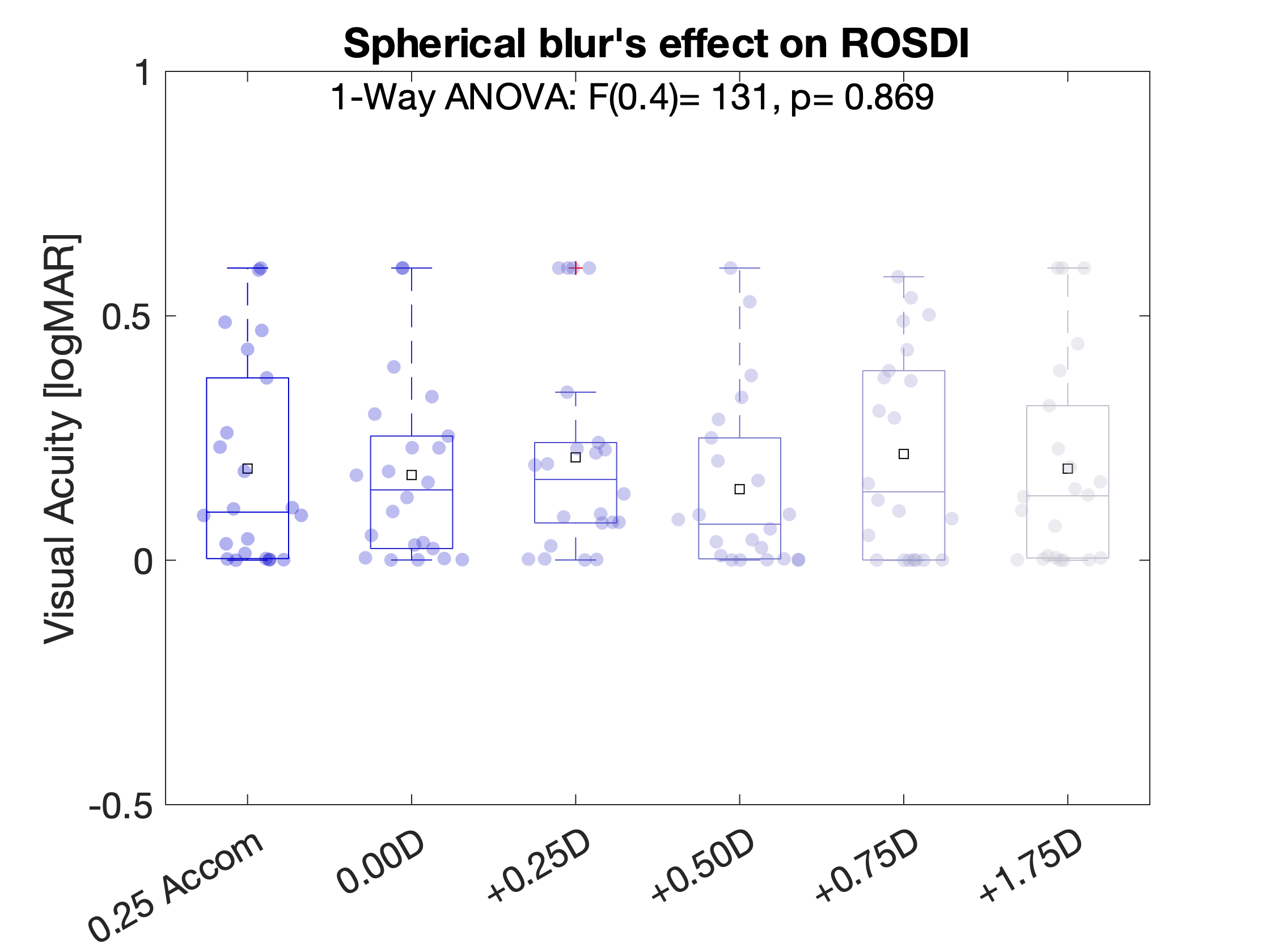

*Figure S6: Effect of spherical blur on the ROSDI (Range Of Stimulus Detectability Improvement). A one-way ANOVA showed no significant effect of spherical blur.*

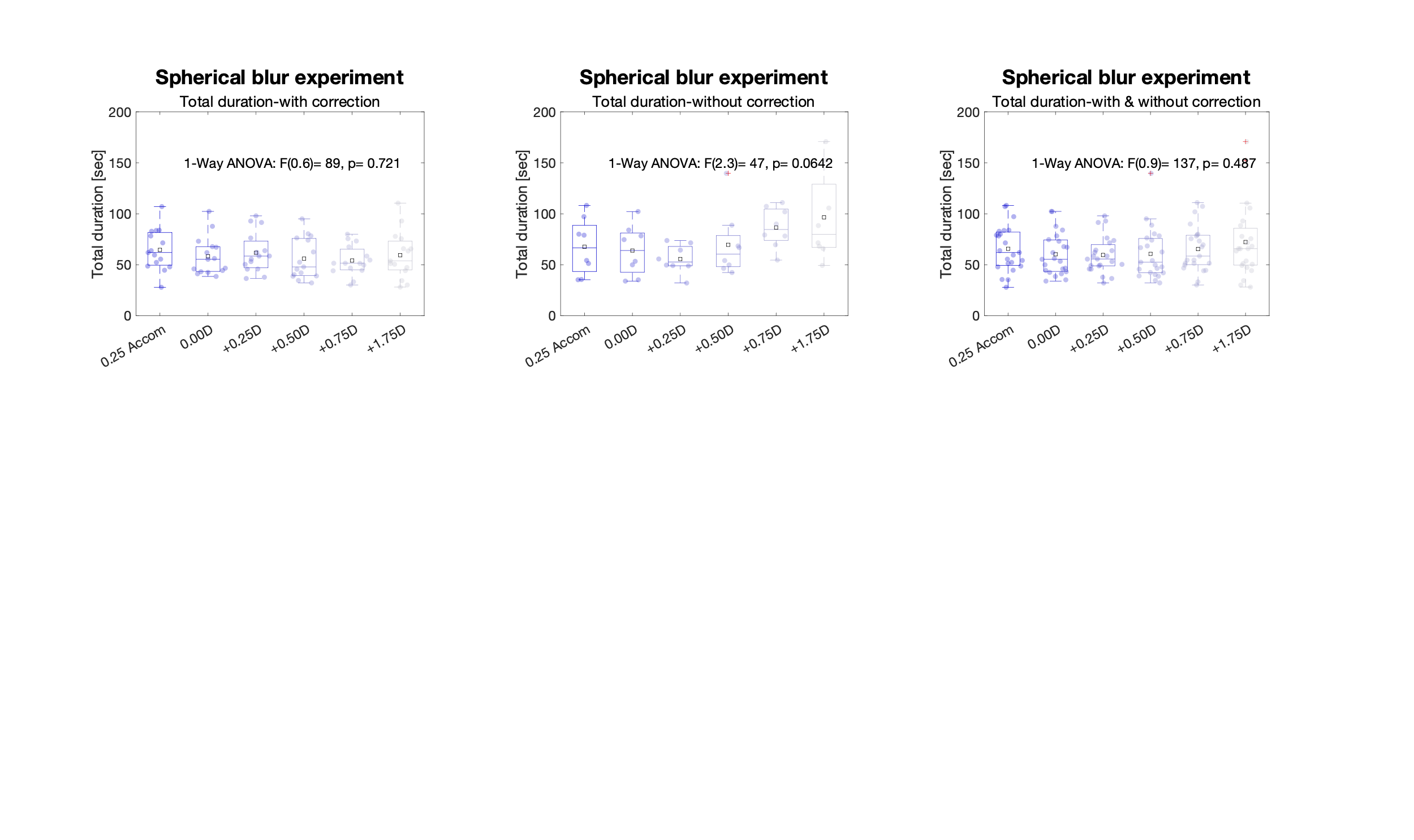

*Figure S7: Effect of spherical blur on AIM-VA assessment time. Y axes show the total test durations (for 2 AIM charts) as a function of spherical blur (x axes) for the group with (left) and without (middle) correction and combined (right). A one-way ANOVAs showed no significant effect of spherical blur on test duration.*

### Experiment 1: Testing AIM’s ability to detect induced astigmatic blur

***Table S2: Planned comparison results for difference between AIM and ETDRS under induced astigmatism using cylindrical lenses. Shown are comparisons between AIM and ETDRS values for each blur condition, including 95% confidence intervals, differences of means between each stimulus type, and p-values for testing for differences.***

| **AIM vs. ETDRS** | **Lower 95%CI** | **Difference means** | **Upper 95%CI** | **P-value** |
| --- | --- | --- | --- | --- |
| **+0.50x0°** | -0.0231830986368668 | 0.121579365704230 | 0.266341830045326 | 0.235700880004789 |
| **+0.50x90°** | 0.0188223958050714 | 0.163584860146168 | 0.308347324487264 | **0.00996661589399167** |
| **+0.50x135°** | -0.0215195880414768 | 0.123242876299620 | 0.268005340640716 | 0.214679885667988 |
| **+1.00x0°** | 0.00618303645400592 | 0.150945500795102 | 0.295707965136199 | **0.0304026933353649** |
| **+1.00x90°** | 0.0481885308959442 | 0.192950995237041 | 0.337713459578137 | **0.000471278407888568** |
| **+1.00x135°** | 0.00784654704939594 | 0.152609011390492 | 0.297371475731589 | **0.0264470545052771** |
| **+2.00x0°** | -0.0165807108620333 | 0.128181753479063 | 0.272944217820160 | 0.159943259657219 |
| **+2.00x90°** | 0.0254247835799050 | 0.170187247921001 | 0.314949712262098 | **0.00529717679539734** |
| **+2.00x135°** | -0.0149172002666433 | 0.129845264074453 | 0.274607728415550 | 0.144039179773353 |

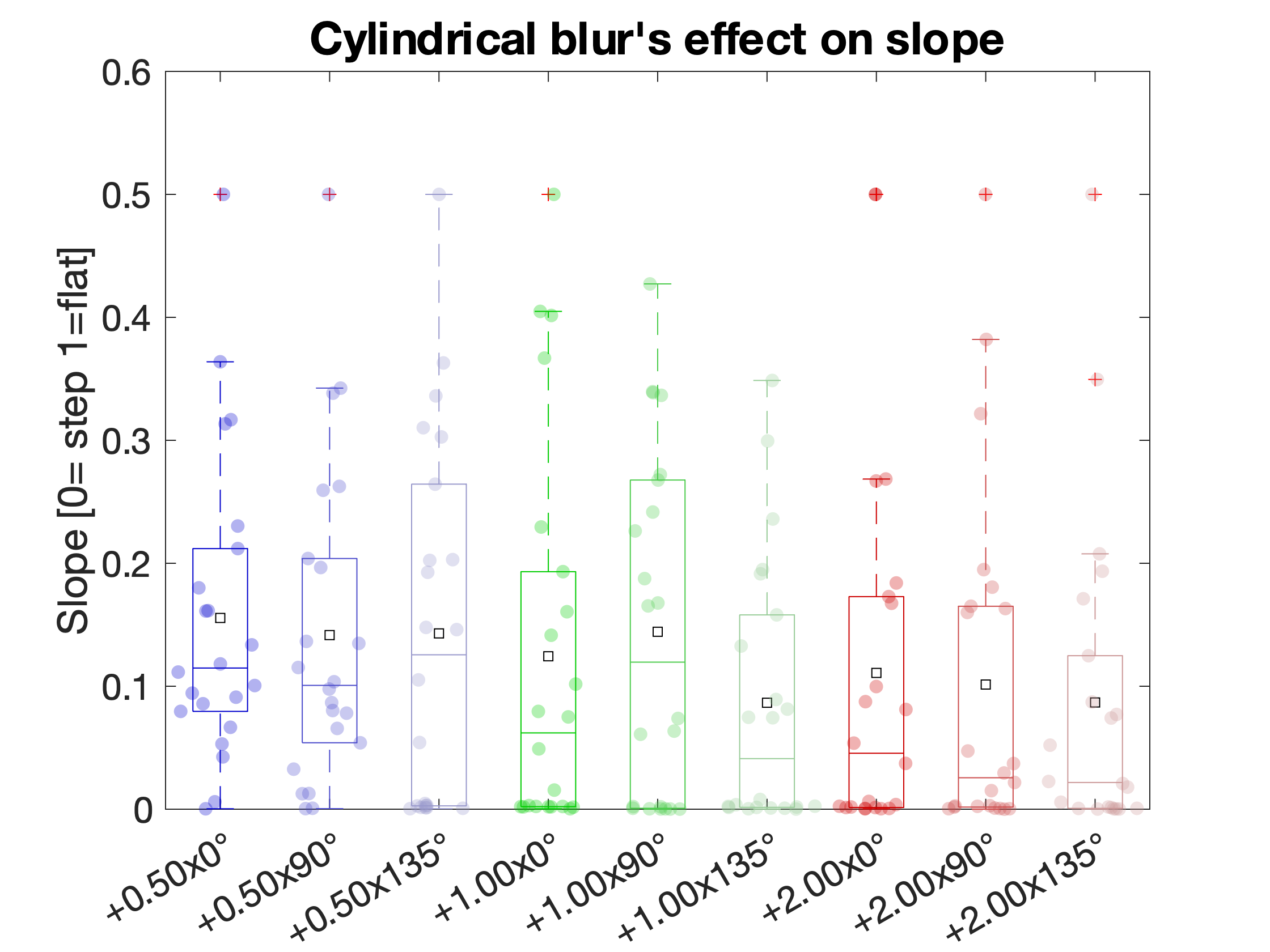

*Figure S8: Comparisons of slope across increasing cylindrical blur levels and axis directions.*

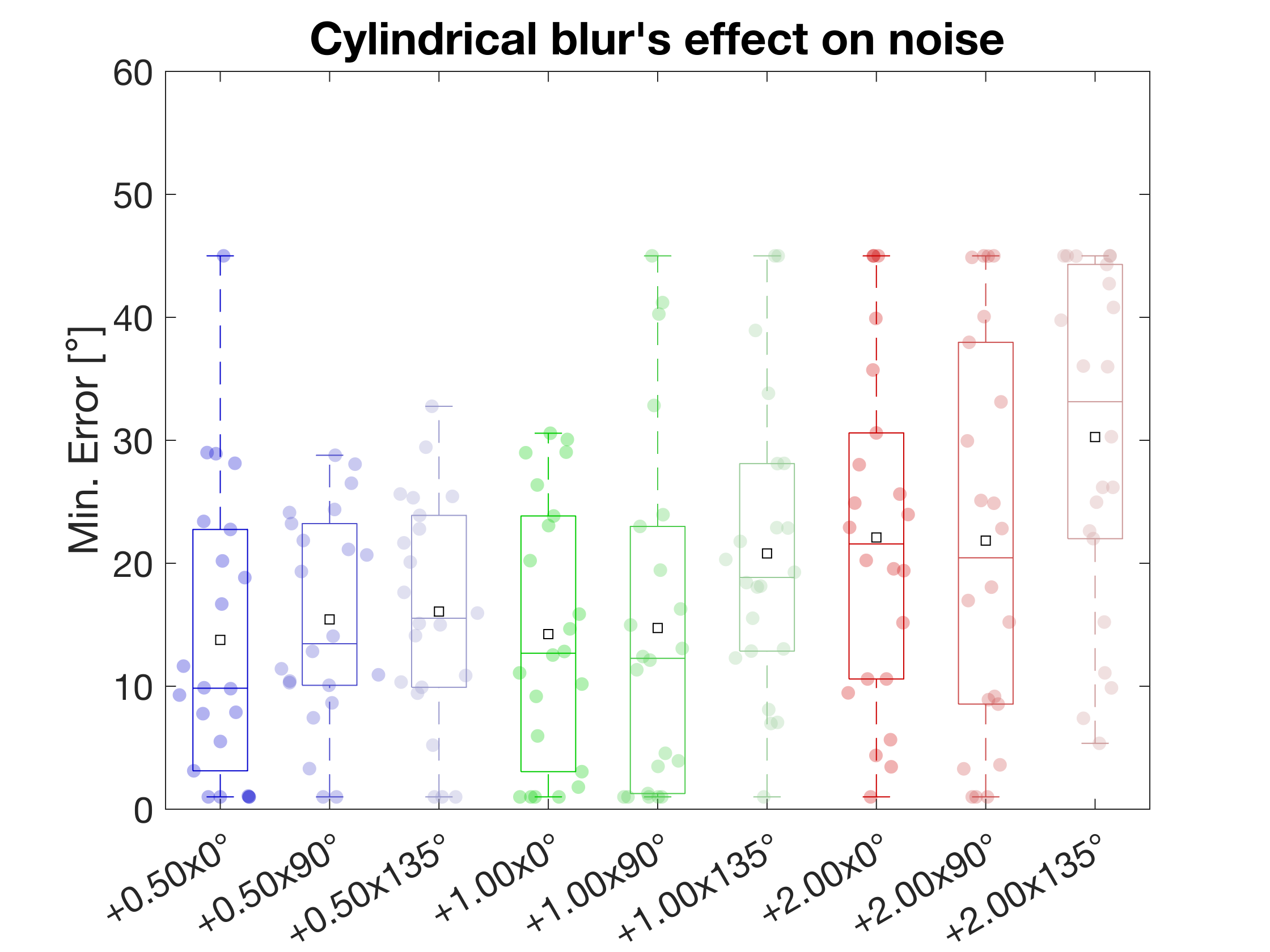

*Figure S9: Comparisons of min. angular error (noise) across increasing cylindrical blur levels and axis directions.*

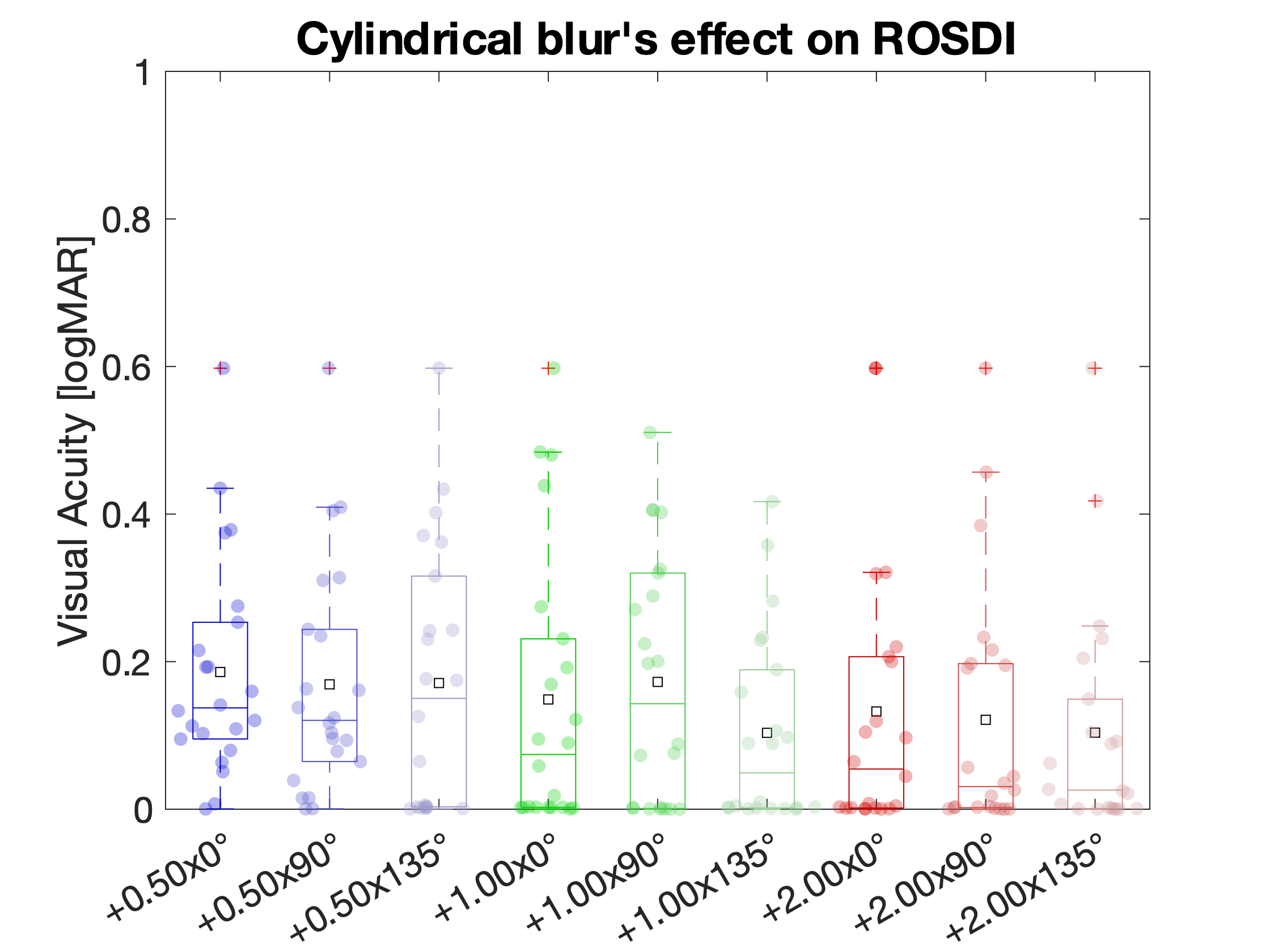

*Figure S10: Comparisons of ROSDI (Range Of Stimulus Detectability Improvement) across increasing cylindrical blur levels and axis directions.*

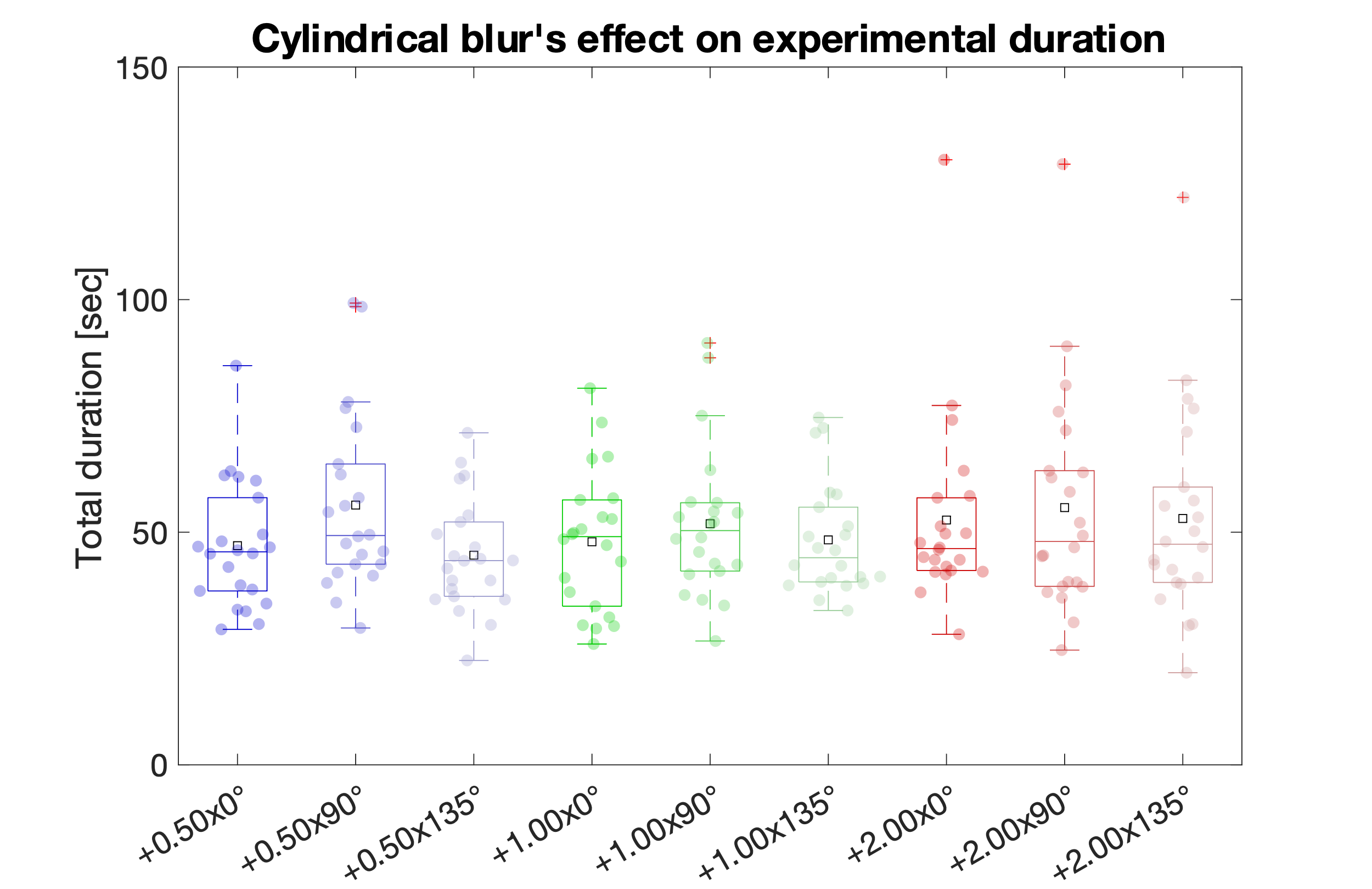

*Figure S11: As Figure S9, except for Cylindrical (astigmatism) blur. Comparisons of experimental duration for AIM across increasing cylindrical blur levels and axis directions. A two-way ANOVA showed no significant effect of blur level [F(2,189) =1.3,p>0.05, η_p_^2^= 0.01] and cylindrical axis orientation [F(2,189)=2.0,p>0.05, η_p_^2^= 0.02] but no interaction [F(4,189)=0.4,p>0.05, η_p_^2^= 0.008].*

#### **Orientation report error analysis using the sinewave model**

In the following, we show the data for the sinewave model’s outputs, namely phase and amplitude of the function.

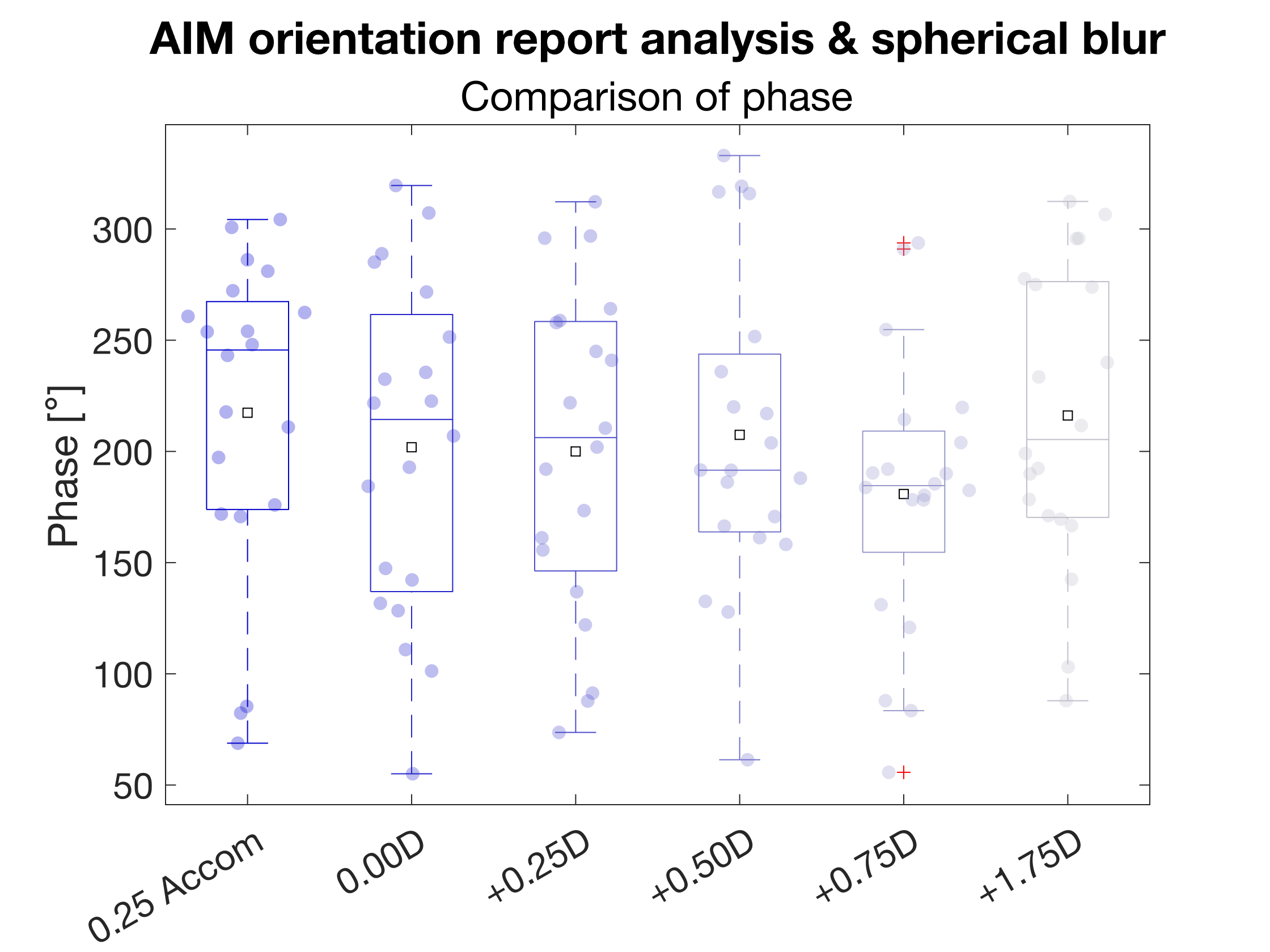

*Figure S12: Results for changes of the phase shifts of the sinewave model due to induced spherical blur.*

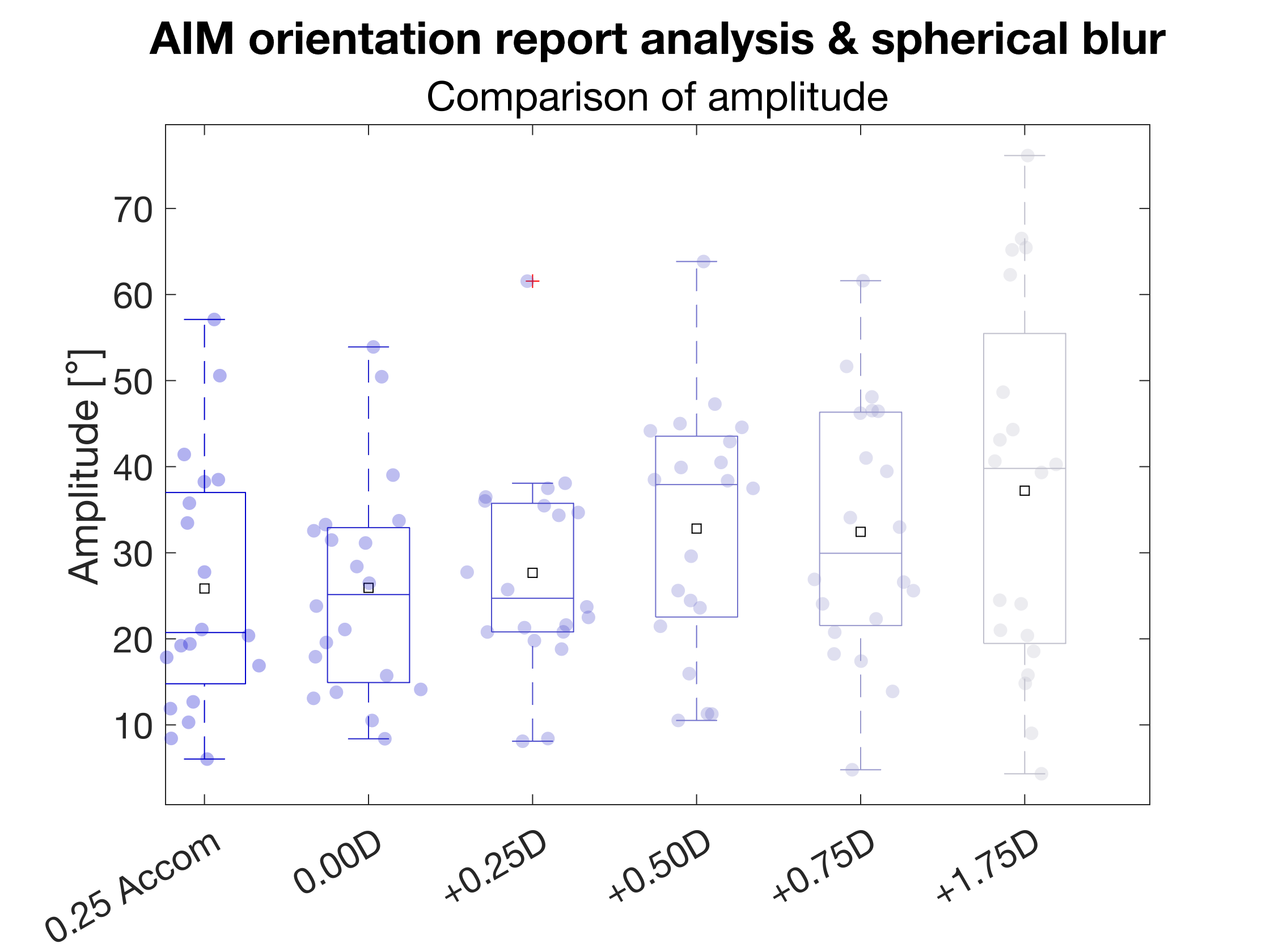

*Figure S13: Results for changes of the amplitude of the sinewave model due to induced spherical blur.*

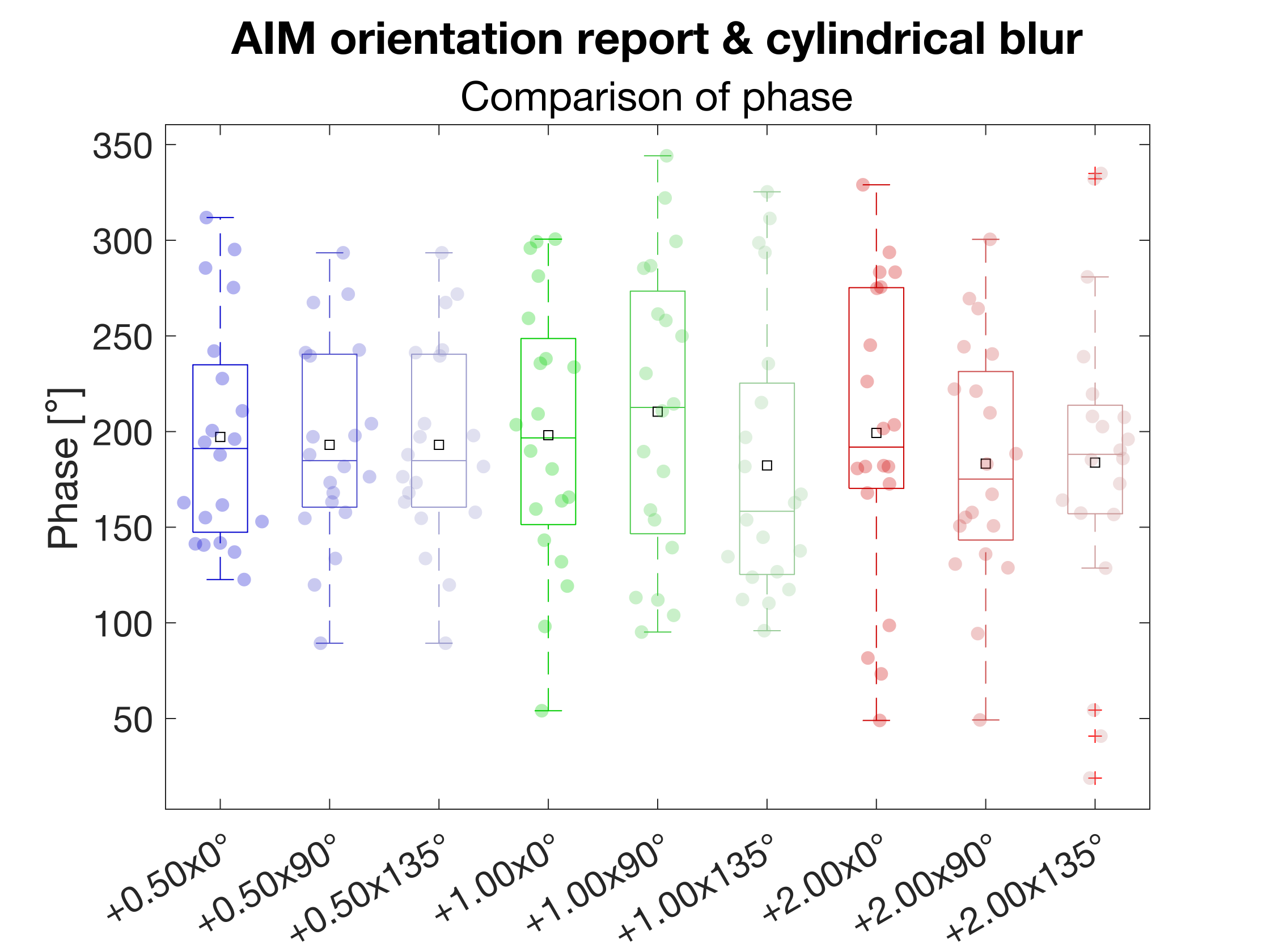

*Figure S14: Results for changes of the phase of the sinewave model due to induced astigmatic blur.*

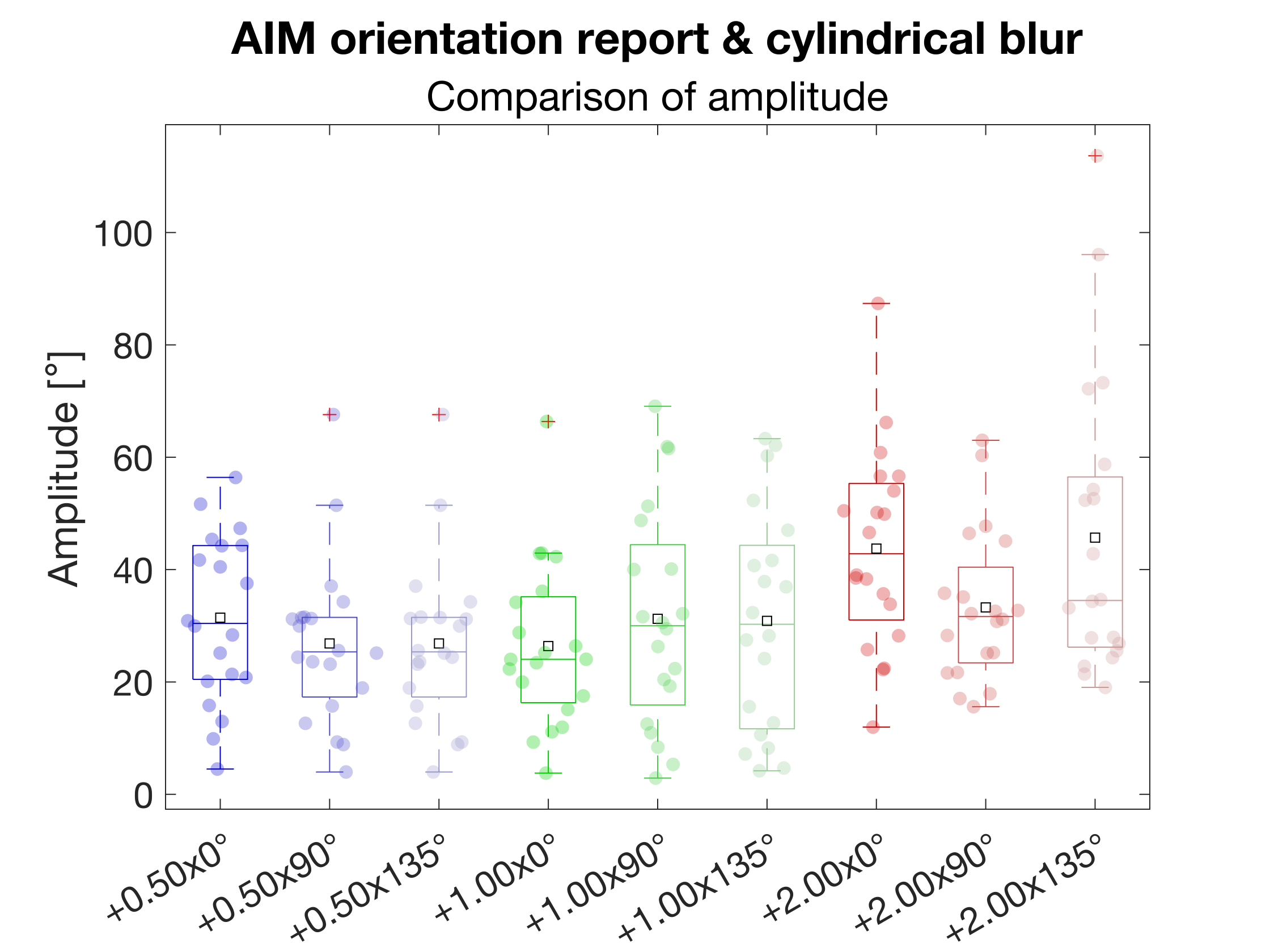

*Figure S15: Results for changes of the amplitude of the sinewave model due to induced astigmatic blur.*
